# Cellular complexity and crosstalk in murine TNF-dependent ileitis: Different fibroblast subsets propel spatially defined ileal inflammation through TNFR1 signalling

**DOI:** 10.1101/2023.09.29.560107

**Authors:** Lida Iliopoulou, Christos Tzaferis, Alejandro Prados, Fani Roumelioti, Vasiliki Koliaraki, George Kollias

## Abstract

Crohn’s disease represents a persistent inflammatory disorder primarily affecting the terminal ileum. Through the application of single-cell RNA sequencing, we unveil the intricate cellular complexities within murine TNF-dependent ileitis, developing in *Tnf*^ΔARE^ mice. Detailed immune cell analysis highlights B cell expansion, T cell effector reprogramming, and macrophage lineage shifts during inflammation. Focusing on stromal cells, we reveal a strong pro-inflammatory character, acquired by all fibroblast subsets, which exhibit complex communication patterns with the infiltrating immune and surrounding stromal cells. Interestingly, we identify that *Tnf*^ΔARE^-induced ileitis is initiated in the lamina propria via TNFR1 pathway activation in villus-associated fibroblasts (Telocytes and Pdgfra^low^ cells). Furthermore, we unveil separate spatial subsets of fibroblasts acting as exclusive responders to TNF, each orchestrating inflammation in different intestinal layers. Additionally, manipulating the *Tnfrsf1a* gene exclusively in fibroblast subsets suggests that inflammation is initiated by telocytes and Pdgfra^low^ cells, while trophocytes drive its progression. This introduces novel evidence of spatial regulation of inflammation by fibroblast subsets, inciting and advancing disease in different layers of the gut. These findings underscore the pivotal role of fibroblasts in the inception and advancement of ileitis, proposing that targeting different fibroblast populations could impede the disease development and chronicity of inflammation.

## Introduction

Inflammatory Bowel Disease (IBD) comprises two primary clinical subtypes, namely Crohn’s disease (CD) and Ulcerative Colitis (UC). Both subtypes involve the development of chronic inflammation in the gastrointestinal tract, but differ in their underlying mechanisms and sites of manifestation.^1,2^ CD primarily affects the terminal ileum, whereas UC is localized to the colon.^2^ Furthermore, in CD, there is frequently an infiltration of immune cells into the deeper layers of the intestinal wall, including the submucosa and muscularis mucosa, often accompanied by the clustering of macrophages within granulomas.^3^

Tumor necrosis factor (TNF) plays a significant pathogenic role in the development and progression of IBD, as evidenced by the successful use of anti-TNF therapy in IBD patients for over 30 years in clinical practice.^4^ *Tnf*^ΔΑRE^ mice, which spontaneously develop CD-like ileitis due to TNF overexpression,^5^ have proven to be a valuable platform for elucidating TNF-dependent mechanisms over the years.^6^

Various intestinal stroma and immune cell types actively participate in the progression of CD.^7,8^ Leveraging the capabilities of single-cell RNA sequencing (scRNA-seq) methodologies, several studies have provided insights into the cellular heterogeneity within the inflamed intestine.^9^ The results have facilitated the discovery of cell subsets that are enriched during the disease, the link between IBD-risk genes and specific cell clusters, the identification of crucial pathogenic pathways, and the correlation of gene signatures with treatment response.^9^

On top of the well-studied hematopoietic cells and epithelial cells, intestinal fibroblasts have emerged as crucial regulators of intestinal inflammatory responses.^8,10^ Different studies have highlighted the appearance of a pro-inflammatory type of fibroblasts during IBD, characterized by the expression of immune-related genes such as *Ccl19*, *Cd74*, and *Il11*.^11,12,13,14^ However, the association of these IBD-related fibroblasts with the homeostatic subsets, as well as their spatial location in the intestine are poorly studied. Moreover, although, the cellular cross-talks between fibroblasts have been examined with more targeted in-silico approaches,^11^ their overall communication patterns during disease have not yet been fully appreciated.

In the present study, we take advantage of the *Tnf*^ΔARE^ CD-ileitis mouse model to generate a single cell atlas that describes the relative cell-abundance alterations, the differentially activated signalling pathways, and the cellular interplays between the immune, and stroma cell compartments during TNF-dependent ileitis. We also provide direct proof that ileitis initiates in the lamina propria through the activation of the TNFR1 pathway in villi associated fibroblasts (Telocytes and Pdgfra^low^ cells). We finally demonstrate, using two complementary fibroblast-specific genetic tools, the requirement of TNFR1 signalling by other fibroblast populations, including the trophocytes for the full progression of ileitis in the deeper intestinal layers. Thus, we pose spatially and functionally distinct fibroblast subsets as the exclusive TNF responder cells that orchestrate inflammation in the different intestinal locations.

## Results

### 1/Intestinal immune and stromal cell atlas of murine TNF-dependent ileitis

To address the cellular complexity of murine TNF-dependent ileitis, we analysed the immune and stromal cells (in total 40.967 cells after filtering and doublet selection process) isolated from the last 6 cm of the ileum from 3-month-old *Tnf* ^ΔARE^ (18778 cells) and *Tnf*^+/+^ (22189 cells) mice with established inflammation in lamina propria expanded in the submucosa and muscularis propria (SFig. 1a). We deliberately excluded epithelial cells from our analysis, as our primary focus was directed towards deciphering the cellular interactions between immune cells and stromal cells within the mucosal environment. To ensure sufficient representation of all the basic cellular compartments, especially given that stromal cells in *Tnf*^ΔARE^ accounted for less than 10% of the total cell population, we sorted myeloid cells (CD45^+^CD11B/CD11C^+^), lymphoid cells (CD45^+^CD11B^−^CD11C^−^), and stroma cells (CD45^−^EPCAM1^−^), mixed them to an equal ratio and performed scRNA-seq. (Fig. 1a, SFig. 1b). Initial analysis on each genotype separately, enabled us to acquire cells from the three primary cellular categories, based on the markers by which they were sorted (Fig. 1b, SFig. 1c).

**Figure 1:**
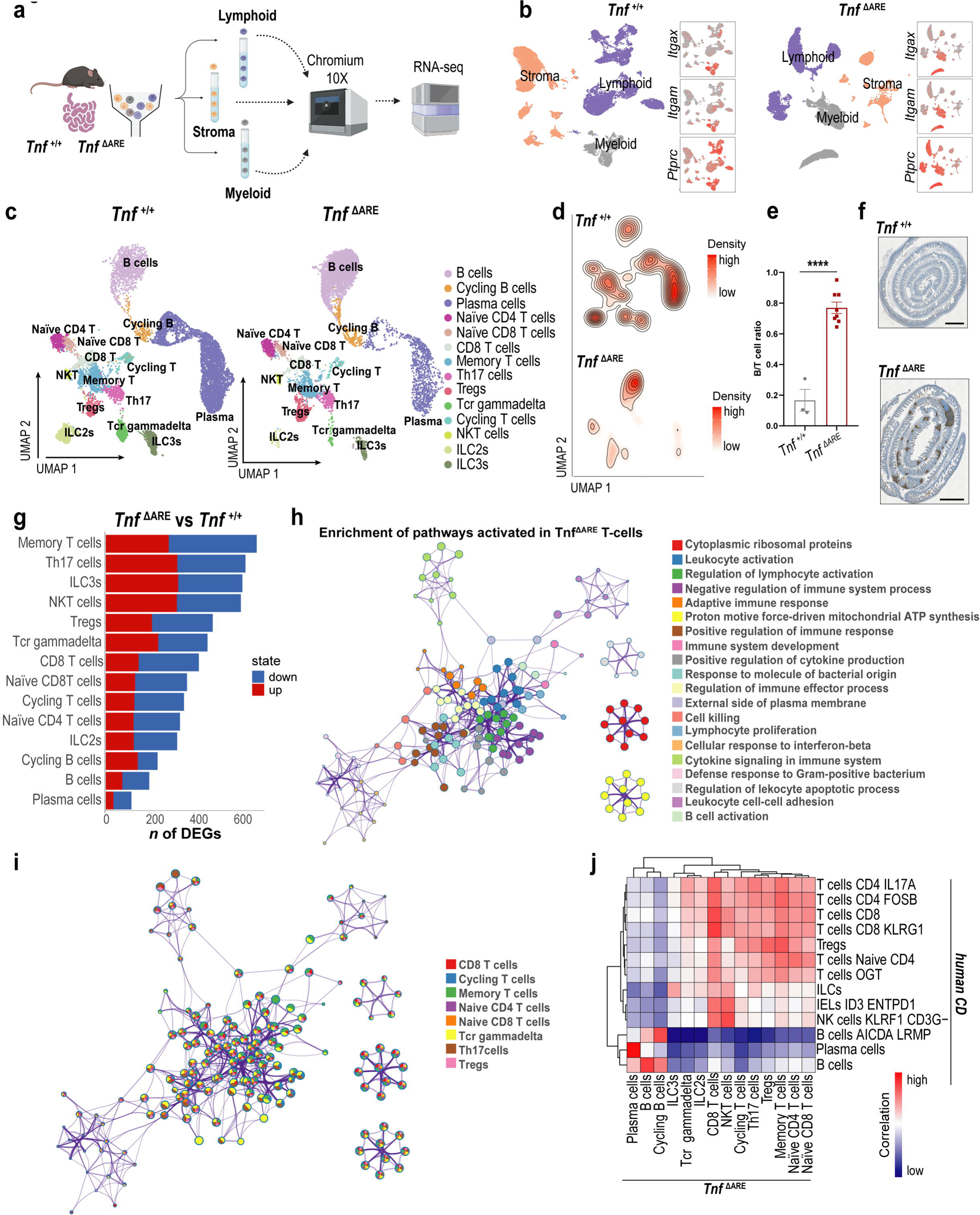
Lymphoid cell atlas of the *Tnf*^ΔΑΡE^ ileum. a/ Schematic representation of the experimental workflow. b/High-quality filtered ileal cells from *Tnf*^+/+^ and *Tnf*^ΔΑΡΕ^ mice (3-months) projected in UMAP space and coloured by cell identity as stroma, lymphoid and myeloid cells. Normalized expression of *Itgam, Itgax and Ptprc* is depicted in feature plots. c/ UMAP projection of integrated *Tnf*^+/+^ and *Tnf*^ΔΑΡΕ^ lymphoid cells. Cells are coloured according to cluster assignment. d/ Contour plot showcasing cell densities in the *Tnf*^+/+^ and *Tnf*^ΔΑΡΕ^ UMAPs. e/ Ileal B to T cell number ratio in the ileum of 3-month old *Tnf*^+/+^ and *Tnf*^ΔΑΡΕ^ mice quantified by flow cytometry, data are mean ± SEM (*n*=3-8 per genotype),two-tailed unpaired Student’s t-test, p.value****<0.001,. f/Immunohistochemistry of B220^+^ cells in ileal swiss rolls from 3-month old *Tnf*^+/+^ and *Tnf*^ΔΑΡΕ^ mice, scale bar=1mm. g/ Barplots representing the number of DEGs resulted from the comparison between *Tnf*^ΔΑΡΕ^ and *Tnf*^+/+^ conditions in all clusters. h,i/ Network representation of the enriched GO biological processes in the *Tn*f^ΔΑRE^ T cells. GO terms have been grouped in clusters by Metascape (h). The depicted GO terms are coloured according to the cluster group identity (i). j/ Heatmap showing the spearman correlation between the most variable genes of *Tnf*^ΔΑΡE^ and CD lymphoid cell subsets.

Initially, we isolated and integrated the lymphoid cells from *Tnf ^+/+^* and *Tnf* ^ΔARE^ mice and further subsetted them into 14 main clusters including B-cells, Plasma cells, ILCs, T cell subsets, NKT cells and cycling lymphocytes (Fig. 1c, SFig. 2a, STab.1). Among the lymphoid cells, we observed a dramatic increase in the relative abundance of B cells (Fig. 1d) in the *Tnf*^ΔARE^ ileum. This increase was further validated through the analysis of the B/T cell ratio, which affirmed the substantial overabundance of B cells in comparison to other lymphocyte populations (Fig. 1e). Indeed, immunohistochemical analysis highlighted the expansion of B cells, distinctly segregated in lymphoid-like structures (Fig. 1f), previously reported to increase in the *Tnf*^ΔARE^ ileum.^5,15^ Although there was a higher relative prevalence of B cells, it is important to highlight that the overall count of intestinal lymphocytes, including IgA^+^ plasma cells, was raised in *Tnf*^ΔARE^ mice (SFig. 2b), indicating a more extensive immune cell infiltration within the intestine.

Τo detect transcriptional changes between the different genotypes, we performed differential expression analysis in the lymphoid cell clusters of *Tnf*^ΔARE^ mice compared to healthy controls (STab. 2). Although the abundance of B cells was largely affected it was notable that the gene expression of newly infiltrated B cells did not exhibit substantial changes (Fig. 1g). Conversely, T cells, particularly memory T cells and Th17 cells, displayed the most significant number of differentially expressed genes (DEGs) (Fig. 1g). Focusing further on the upregulated genes on T cell subsets, we used the Metsacape portal to infer molecular pathways, gene ontology (GO) terms and associations between them, enriched in Tnf*^ΔARE^*cells compared to heathy controls (STab. 3). Prominent biological functions governing T cell activation, such as the regulation of immune effector processes, lymphocyte proliferation, leukocyte cell-cell adhesion, and cellular response to interferon-beta, were markedly enriched within T cell clusters (Fig. 1h), including memory, TCRγδ, CD8, and Th17 cells (Fig. 1i). To identify the similarities in the lymphoid cells between murine and human ileitis, we performed correlation analysis between the *Tnf^ΔARE^* and a CD lymphoid dataset.^16^ B cells, Plasma cells and ILC3s presented a high degree of correlation between diseased organisms (Fig. 1j). The T cell populations displayed robust correlations with a mixed population of human diseased T cells, with particularly strong and specific the correlation observed between mouse CD8+ T cells and human CD8 and CD8+ KLRG1+ cells, as well as between mouse NKT cells and human NK cells and IELs ID3/ENTPD1, in addition to the correlation between mouse and human Tregs, and mouse and human naïve T cells (Fig. 1j). This alignment in gene expression patterns between lymphoid cells in murine and human CD ileum is indicative of a noteworthy degree of transcriptional similarity during disease.

Collectively, these observations propose the expansion of B cells over the rest of lymphoid cells in the *Tnf*^ΔΑΡE^ ileum and the transcriptional remodelling of T cell subsets towards acquiring more effector and memory functions.

### 2/A monocyte to LYZ1+ of macrophage lineage dominates the *Tnf*^ΔARE^ ileum

We next sought to examine the heterogeneity of myeloid cells that infiltrate the inflamed murine ileum. After clustering them, we annotated 9 distinct subsets (SFig 3a, STab.1); including granulocytes, monocytes, 3 types of macrophages: resident, activated macrophages, and macrophages with an intermediate phenotype between monocytes and mature macrophages, 4 clusters of dendritic cells: conventional type 1 DC (cDC1), conventional type 2 DC (cDC2a and cDC2b) and plasmacytoid dendritic cells (pDCs) (Fig. 2a, Fig. 2b). While in healthy mice, we barely detected granulocytes, they got massively increased in inflamed state, together with monocytes, activated macrophages and the cDC2a subset of DCs (Fig.2c).

**Figure 2:**
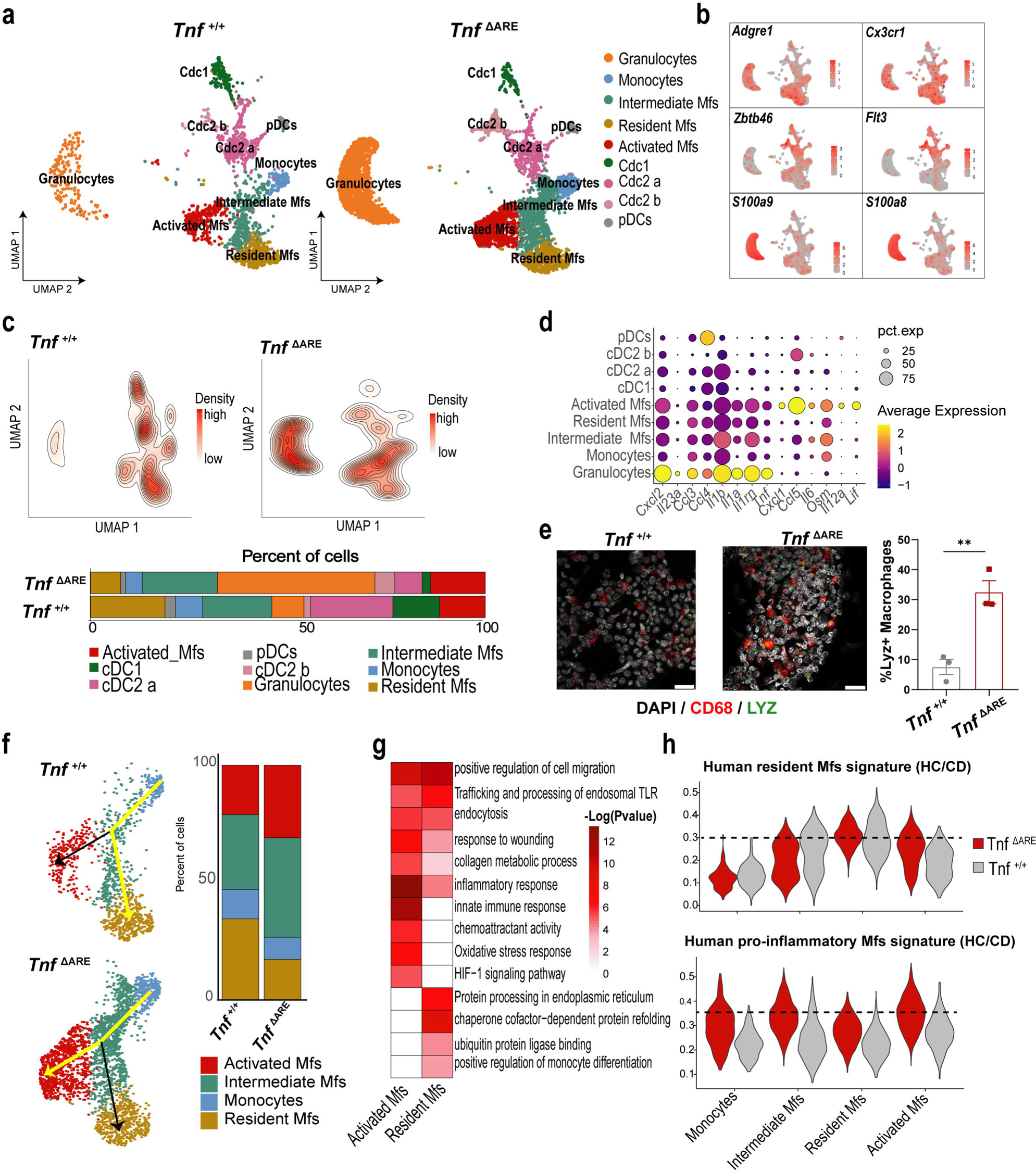
Transcriptome heterogeneity of ileal myeloid cells during inflammation. a/ UMAP-plot, depicting subclusters of myeloid cells (CD45^+^CD11B^+^CD11c^+^) from the ileum of *Tnf*^+/+^ and *Tnf*^ΔΑΡΕ^ mice. b/ Feature-plots displaying the normalised expression of gene markers associated to basic myeloid populations. c/ Contour plots and barplots depicting the differential abundance of myeloid cell clusters per genotype. d/Dotplot of the average normalised gene expression of pro-inflammatory cytokines and chemokines per cluster. e/Immunofluorescent detection of CD68 and LYZ1 in cytospin preparations of ileal single cell suspensions from 3-month old *Tnf*^+/+^ and *Tnf*^ΔΑΡΕ^ and quantification data of double CD68^+^LYZ1^+^ macrophages; data are mean ± SEM (*n*=3 per genotype), two-tailed unpaired Student’s t-test,**p.value < 0.01 f/ Pseudotime-trajectory analysis depicting the two distinct lineages of myeloid cells color coded according to the final state (yellow lines indicate which differentiation path is enriched in each genotype) and barplots representing the different cell subsets of the trajectory across genotypes. g/Heatmap representing shared and unique pathways enriched in resident and activated macrophages identified using Metascape. h/Violin-plots showing activity scores of the CD-related human resident and pro-inflammatory macrophages gene signatures in the mouse myeloid clusters.

By mapping the expression of basic pro-inflammatory cytokines and chemokines in the myeloid clusters, we identified granulocytes and activated macrophages as the main cellular sources of distinct immune-related signals (Fig 2d); Granulocytes highly express *Ccl3*, *Ccl4*, *Cxcl12* and important effector cytokines such as *Tnf* and members of the interleukin-1 (IL-1) family (*Il1a, Il1b, Il1rn*), while activated macrophages predominantly expressed *Ccl5*, *Cxcl1*, and members of the IL-6 family (*Osm*, *Il6*, and *Lif).* Alongside these pro-inflammatory genes, activated macrophages were defined by a set of unique markers including *Lyz1, Gpnmb* and *Saa3* (SFig. 3a). *Lyz1* is considered a specific marker for Paneth cells.^17^ Indeed, we identified an increase of LYZ1^+^ macrophages in the inflamed ileum of *Tnf*^ΔΑRE^ mice (Fig. 2e). Trajectory analysis of macrophage populations using the Slingshot R package, uncovered the existence of two monocyte-derived lineages that share as a transitional point the intermediate macrophages and are characterized by two distinct final differentiation states, the resident and the activated macrophages (Fig. 2f). The lineage that leads to the resident macrophages was enriched in *Tnf*^+/+^ mice, while the activated macrophage lineage was over-represented in the *Tnf*^ΔΑRE^ ileum (Fig. 2f, SFig. 3b), supporting the notion that monocyte-derived, inflammation-associated macrophages are originally separated from the resident intestinal macrophages. In line with this, applying the metascape analysis in the markers genes of resident and activated macrophages, highlighted unique molecular pathways (STab.3). Resident macrophages were characterized by enhanced activity in pathways related to antigen presentation, whereas activated macrophages showed a dominant pro-inflammatory response, accompanied by oxidative stress (ROS production), hypoxia, and collagen metabolism (Fig. 2g). Indeed, activated macrophages presented high expression of genes related to extracellular matrix (ECM) degradation (*Mmp9, Mmp12, and Mmp14*) (SFig. 3c) previously been associated to the granuloma-related macrophages.^18^ Immunofluorescence analysis Immunofluorescence analysis confirmed the presence of macrophages within granuloma-like structures, preferentially localized in the submucosa of diseased mice (SFig. 3d).

Previous scRNA-seq analysis in the ileum of human CD patients stressed the existence of two macrophage subsets, attributed as resident and pro-inflammatory macrophages.^11^ Mapping their gene signatures (STab. 4, SFig. 3e) to mouse myeloid cells suggested a strong association of human and mouse resident macrophages and a substantial correspondence of human CD pro-inflammatory macrophages with the activated and intermediate *Tnf*^ΔΑΡE^ macrophage subsets (Fig. 2h).

These findings highlight the similarities shared in the transcriptional profile of disease-associated ileal macrophages in human and murine CD-ileitis.

### 3/Compositional and transcriptional remodelling of stroma cells during ileitis

Following the analysis of immune cells, we aimed to characterize in detail the stromal cells during intestinal inflammation. Integration and unsupervised clustering revealed 14 cellular clusters. Detailed examination of representative markers identified clusters of endothelial cells (lymphatic, vein, and capillary), fibroblasts (telocytes, trophocytes, Pdgfra^low^, FRCs, FDCs/MRCs), smooth muscle cells (SMCs), glial cells, mesothelial cells, interstitial cells of Cajal and pericytes (Fig. 3a, SFig. 4a). We next assessed the relative abundance of stromal cell clusters in *Tnf*^+/+^ and *Tnf*^ΔΑRE^ mice. We detected a marked reduction in fibroblast subsets (telocytes, trophocytes, Pdgfra^low^) and a relative expansion of lymphatic endothelial cells (LECs) and fibroblastic reticular cells (FRC) populations (FRC1&FRC2) (Fig. 3b, SFig4b). The overrepresentation of FRCs was expected in the areas where TLOs developed.^15,19^ Indeed, we validated the decline in the overall population of fibroblasts, both as number and percentage in 3-month old *Tnf*^ΔΑRE^ mice (Fig. 3b, SFig. 4c), suggesting that the composition of stromal cells is affected, possibly due to the shortening and blunting of the inflamed villi in the ileal mucosa. Following the dominant changes in their relative abundance, fibroblast subsets exhibited the highest number of DEGs (telocytes: 998, Pdgfra^low^: 809, trophocytes: 669) together with LECs (781) and SMCs (823) (Fig. 3c). In contrast, FRCs showed minimal gene expression alteration compared to wild-type subsets (FRC1: 158, FRC2: 152) (Fig. 3c). Enrichment analysis in the upregulated genes of stromal subsets, highlighted a strong pro-inflammatory signature acquired primarily by the telocytes, trophocytes, Pdgfra^low^, capillary ECs, and SMCs (Fig 3d). This was characterized by pathways associated with cytokine response (TNF signaling pathway, response to interleukin-1, response to type II interferon), innate immune response (NOD-like receptor signaling pathway, response to lipopolysaccharide, regulation of chemotaxis), and antigen presentation (antigen processing and presentation). Fibroblast together with endothelial cell clusters upregulated also genes related to vasculogenesis and new blood vessel cell formation (Fig. 3d).

**Figure 3.**
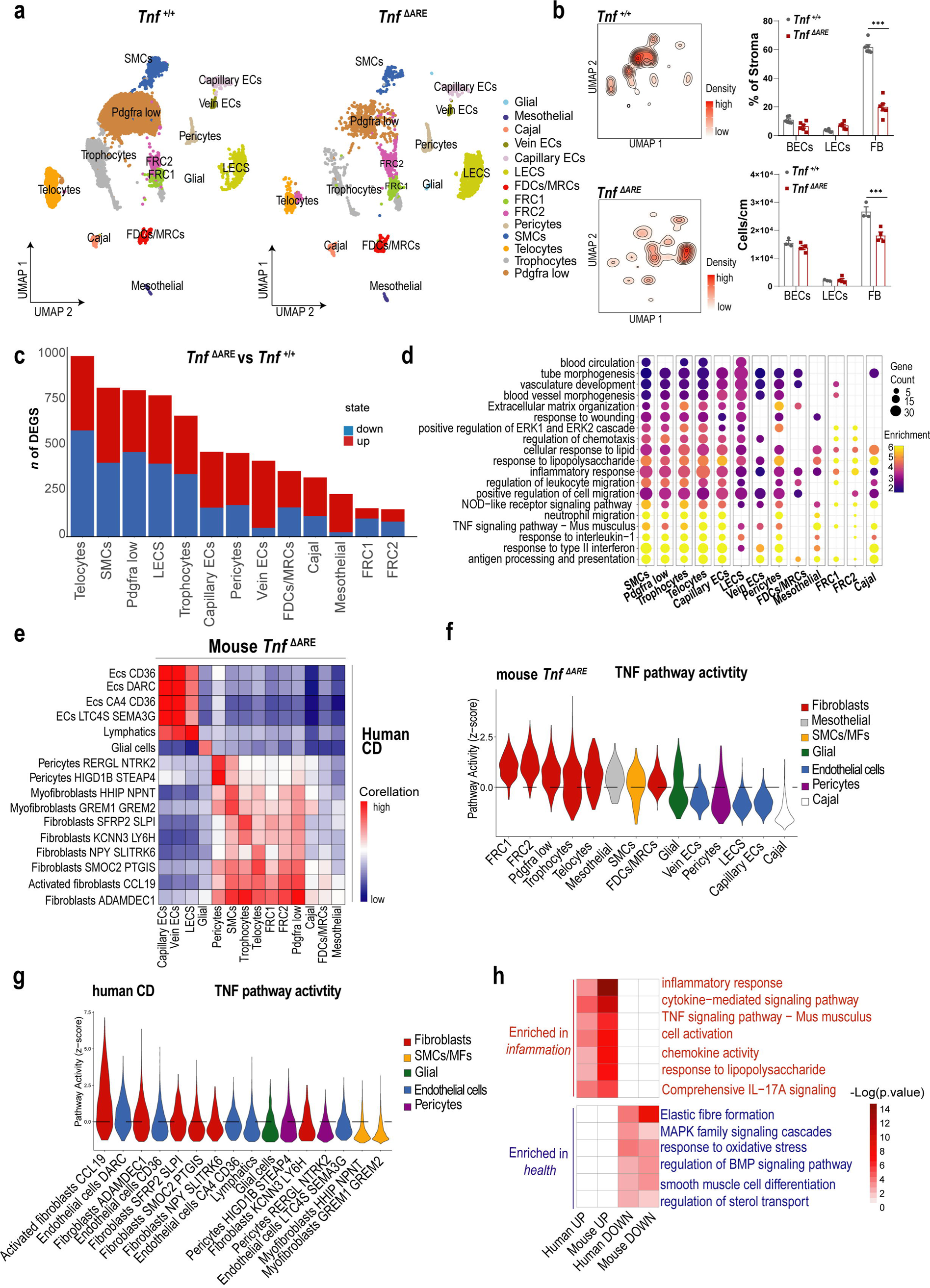
Fibroblast subsets are highly affected compositionally and transcriptionally upon inflammation in the ileum. a/Integration, UMAP projection, and cell annotation of ileal *Tnf*^+/+^ and *Tnf*^ΔARE^ stromal cells (CD45^−^EPCAM^−^). b/ Contour plot representing the cell abundancies of stromal cell subsets projected in the UMAP space. Flow cytometry analysis of the absolute number and percentage (in the CD45^−^EPCAM^−^ cells) of endothelial cells and fibroblasts in the ileum of 3-month-old mice with the indicated genotypes. Each dot represents sample from one mouse, data are ± SEM, n=3-5, Ordinary 2-way Anova, ***p.value<0.001. c/ Barplots showing the number of DEGs in the stromal clusters (*Tnf*^ΔARE^ versus *Tnf*^+/+^). d/Dotplot depicting the enrichment score of biological processes/pathways in the upregulated genes of *Tnf*^ΔARE^ clusters using Metascape. e/ Correlation analysis between *Tnf*^ΔARE^ and CD stromal cells. f,g/TNF pathway activity calculated through *PROGENy* algorithm in *Tnf*^ΔARE^ (f) and human CD (g) stromal cells. h/ Heatmap displaying shared enriched and downregulated pathways between mouse and human diseased fibroblast superclusters.

To examine the relevance of our data with human CD, we performed correlation analysis with a previously reported stromal cell dataset including cells from CD inflamed ileum, uninvolved areas, and healthy specimens.^16^ There was a strong and specific correlation between mouse and human diseased endothelial subtypes, glial cells, pericytes, and myofibroblasts/SMCs (Fig. 3e). Regarding fibroblast subtypes, human CD fibroblasts SFRP2 SLPI and KCNN3 LY6H were mostly correlated with *Tnf*^ΔΑRE^ trophocytes, while CD fibroblasts NPY SLITRK6 were best correlated with *Tnf*^ΔΑRE^ telocytes (Fig. 3e). Among fibroblasts, ADAMDEC1, the activated fibroblasts ADAMDEC1 CCL19 and the SMOC2 PTGIS clusters were equally similar to all mouse fibroblast subsets, suggesting that these human clusters represent a mixed population of cells corresponding to mouse subsets (Fig. 3e). Notably, the broad correlation of the activated ADAMDEC1 CCL19 fibroblasts with the *Tnf*^ΔΑRE^ subsets proposes that CD-associated cell subsets (like the CCL19+ fibroblasts) may be reflected in various mouse fibroblasts populations and may not represent nessecerily unique disease emerging clusters.

Since TNF is the driver pathway for the development of ileitis, we inferred the TNF pathway activity score on the stromal cell subsets using *PROGENy*.^20^ Surprisingly, *Tnf*^ΔΑRE^ fibroblasts subsets (FRC1, FRC2, telocytes, Pdgfra^low^, and trophocytes) presented the highest activation of TNF pathway (Fig. 3f). Interestingly, human CD fibroblasts and especially the disease associated ADAMDEC1 CCL19 cluster, were also having an enriched TNF pathway activity together with two subtypes of endothelial cells (Fig. 3g). Supercluster generation in human and mouse datasets (STab. 5) and subsequent differential gene expression and pathway enrichment analysis in CD versus healthy fibroblasts in both mouse and human unveiled common pathways enriched or downregulated in disease (Fig. 3h). Among the shared upregulated biological processes, all related to immune-mediated functions, we stress the upregulation of response to TNF which was evident in both human and mouse diseased fibroblasts as opposed to healthy fibroblast clusters (Fig. 3h).

Overall, our analysis in stromal cell subsets reveal that despite the reduced number of fibroblasts subsets during ileal inflammation, their activation is significantly skewed towards a pro-inflammatory phenotype. Importantly, all fibroblast subsets acquire similar properties, implying that each of them plays a substantial role in the development of the disease. In addition, this activation pattern shares striking resemblances with human CD subtypes, notably featuring pronounced TNF pathway activation.

### 4/Fibroblast communication patterns in the *Tnf*^ΔΑRE^ ileum

To infer the cellular connectome in the diseased and healthy ileum, we performed comparative analysis using *CellChat*.^21^ To generate a more simplified communication network, we categorised the cells into superclusters (STab. 6) and visualized the number of interaction signals (Fig. 4a, SFig. 5a). Surprisingly, among the most evident changes in the *Tnf*^ΔΑRE^ mice was the enhanced paracrine and autocrine fibroblast signals (Fig. 4a). Comparing the strength of outgoing and incoming interactions in a two-dimensional space, facilitated the identification of three cellular hubs that display noteworthy variations in their ability to send or receive signals (Fig. 4b). Class I includes cell subsets characterized by relatively low levels of both incoming and outgoing signals, with the exception of pDCs. Class II incorporates cells demonstrating moderate strength in receiving and sending signals, while Class III encompasses cell clusters displaying high signal transmission and medium receiving capacity. Significantly, in the *Tnf*^ΔΑRE^ mice, all fibroblast subsets and SMCs were prominently represented within the Class III hub, indicating their role as quantitavely primary cellular sources of signals. Conversely, in contrast to healthy mice, the Class II hub in *Tnf*^ΔARE^ mice comprised endothelial cell types, pericytes, mesothelial cells, and activated macrophages, all exhibiting relatively balanced incoming and outgoing capacities (Fig. 4b).

**Figure 4.**
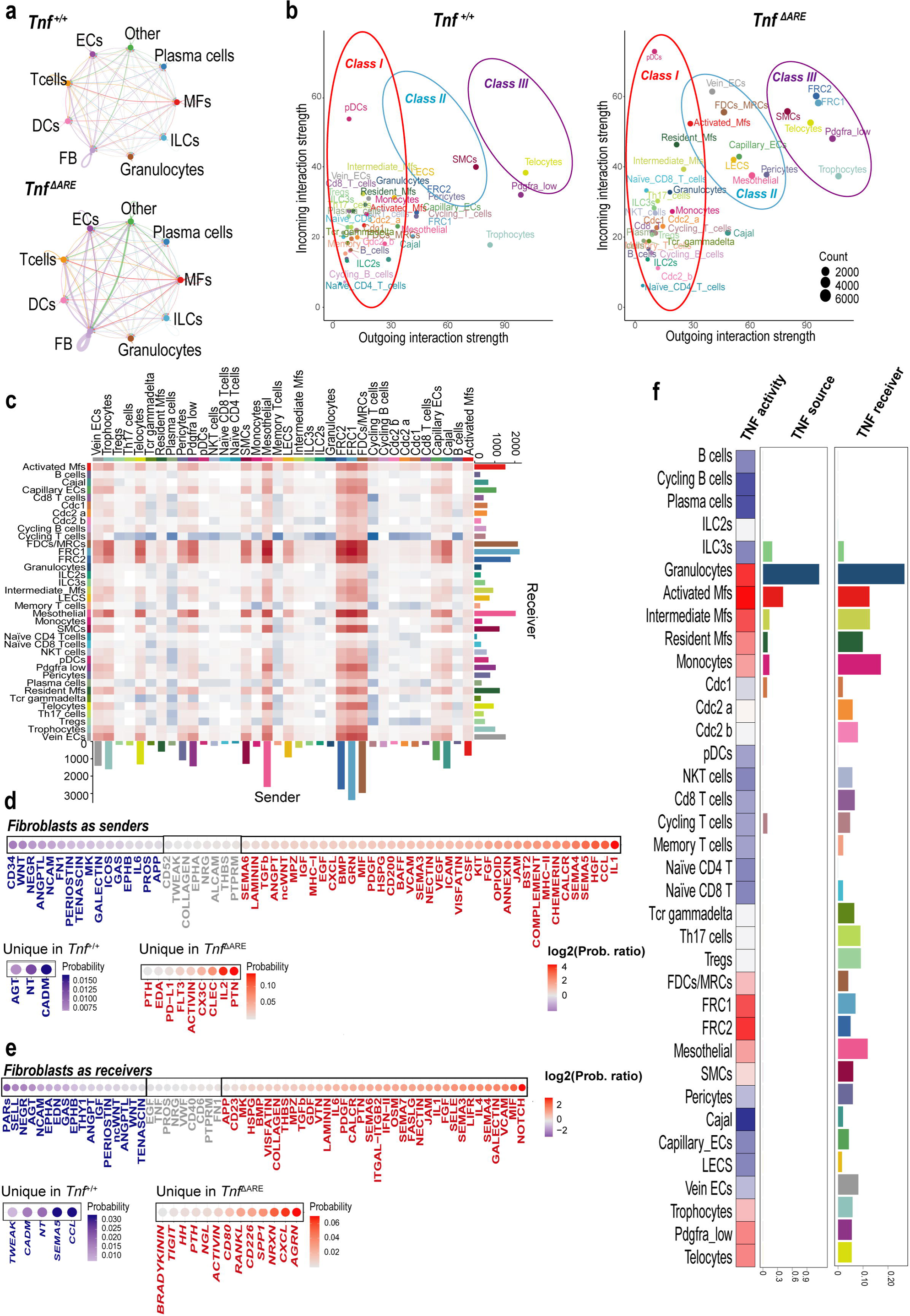
Cell-cell communication analysis reveals active signalling pathways in ileal homeostasis and inflammation. a/Signalling network between cell subsets organized in broad cell categories in *Tnf*^+/+^ and *Tnf*^ΔΑΡΕ^ (3-months) conditions. b/Scatter plot showcasing the differences in incoming and outgoing interactions of each cluster between conditions. c/ Heatmap depicting the differential number of interactions per cluster among the two conditions. The barplots show the aggregated communication probability when a cell cluster is considered as sender (bottom) or as receiver (right). d,e/Dotplots of shared and sample specific signalling pathways when fibroblasts are set as senders (d) or receivers (e). In the first case the dot color denotes the log ratio of the communication probabilities, while in the second the communication probability of the specific sample. f/Heatmap and barplots for the TNF pathway. In the heatmap the PROGENy activity scores are shown per cluster, while in the barplots the cellchat communication probability is depicted again for the TNF signalling pathway.

Next, we visualized the differential number of interactions between the two datasets, including all the cell cluster pairs (Fig. 4c). Interestingly, notable modifications were observed in the paracrine and autocrine signaling patterns specifically within the FRC and FDCs/MRC populations (SFig. 6, SFig. 7). These subsets of fibroblasts primarily reside in the Peyer’s Patches (PPs) and TLOs, where they play a crucial role in supporting the structural integrity and functionality of these lymphoid structures.^22^ To further investigate the necessity of PPs and TLOs in the progression of ileitis, we impeded their formation in *Tnf*^ΔΑRE^ mice by deleting the *Ltbr* gene, which is crucial for the development of peripheral lymphoid organs. As a result, *Ltbr*^KO^ *Tnf*^ΔΑRE^ mice completely lacked PPs and TLOs (SFig. 8a). However, the severity of ileitis remained unaffected by the absence of these lymphoid organs in the ileum, indicating that they are dispensable for the initiation and progression of intestinal inflammation (SFig. 8b). This implies that although FRCs/FDCs/MRCs exhibit an increased number of outgoing and incoming signals, which may be important for the formation of ectopic TLOs in the ileum of *Tnf*^ΔARE^ mice, they are not actively involved in the propagation of inflammation.

Thus, we focused in the communication pattern of non-FRC fibroblasts (trophocytes, telocytes, Pdgfra^low^) that exhibited an increased ability to transmit and receive signals during ileitis. By calculating the differential aggregated probability score between diseased and healthy mice, we observed signals originating from (Fig. 4d) or targeting (Fig. 4e) fibroblasts that were enriched in either *Tnf*^ΔΑRE^ or wild-type mice (STab.7). Examples of eliminated fibroblast-derived signals during ileitis include the expression of CD34 anD WNT ligands, related among other to epithelial cell homeostasis and NCAM ligands that are related to neuronal functions (Fig. 4d). On the other hand, in the upregulated ligands, there was an augmented expression of chemotactic molecules (such as CCLs, CXCLs, SEMA3/4/5/6, and CHEMERIN), adhesion mediators (like ICAM), and pro-inflammatory activators of immune effector cells (including COMPLEMENT, IL1, IL2, FLT3, BAFF, and MHCI/II) (Fig. 4d). Among the fibroblast-targeting signaling pathways, we observed an upregulation of ECM interactions (COLLAGEN, LAMININ) and pro-inflammatory cytokines, including IL-1, type II interferons and STAT3 activators (IL6, OSM, LIFR) (Fig. 4e), all of them reported to be implicated in anti-TNF non-response mechanisms.^23,24,25^

The calculation of communication probabilities relies on the evaluation of ligand and receptor expression within cells belonging to particular clusters.^21^ Interestingly focusing in non-FRC fibroblast clusters, we observe no significant upregulation of *Tnfr1,* and a modest increase in *Tnfr2* expression during ileitis (SFig. 5b), explaining the similar communication propability of TNF signaling between *Tnf*^+/+^ and *Tnf*^ΔΑRE^ mice (Fig. 4e). However, there was clear evidence of TNF overexpression in granulocytes, activated macrophages, and to a lesser extent, in other myeloid cells (Fig. 4f, SFig. 5b). In addition, applying the *PROGENy* algorithm in all *Tnf*^ΔΑRE^ subsets revealed the activation of TNF pathway (incorporating molecular mediators and downstream target genes) primarily in myeloid cells and secondary to fibroblast subpopulations (including telocytes, trophocytes and Pdgfra^low^) (Fig. 4f).

In summary, among all cell types in the ileum (excluding epithelial cells), fibroblast subsets exhibit the highest propability of acting as sender cell types during the disease, driving inflammation through different chemotactic and immune-activating signals. In parallel, they function as receivers of multiple activating stimuli, including TNF, the primary cytokine implicated in TNF-dependent ileitis as well as other signals involved in non-response to anti-TNF mechanisms during disease.

### 5/Differential targeting of spatially organized fibroblast subpopulations in the inflamed ileum

Due to the striking transcriptional activation of fibroblast subsets, we aimed to identify their spatial organization during the development of ileitis. We conducted disease evaluation on *Tnf*^ΔΑRE^ mice at different stages of their development (Fig. 5a). At 3 weeks of age, histological assessment revealed an absence of inflammatory infiltrates. However, at 8 weeks, inflammation in the lamina propria had initiated, accompanied by mild infiltration in the submucosal layer. As the mice reached 3 and 6 months of age, the submucosal layer underwent massive enlargement, leading to an increased submucosal thickness. Notably, in this area, fibroblasts showed a specific expansion according to their spatial distribution. Additionally, the muscularis mucosa displayed mild infiltration at a later disease stage (3 months) and became more established by 6 months of age (Fig. 5a).

**Figure 5.**
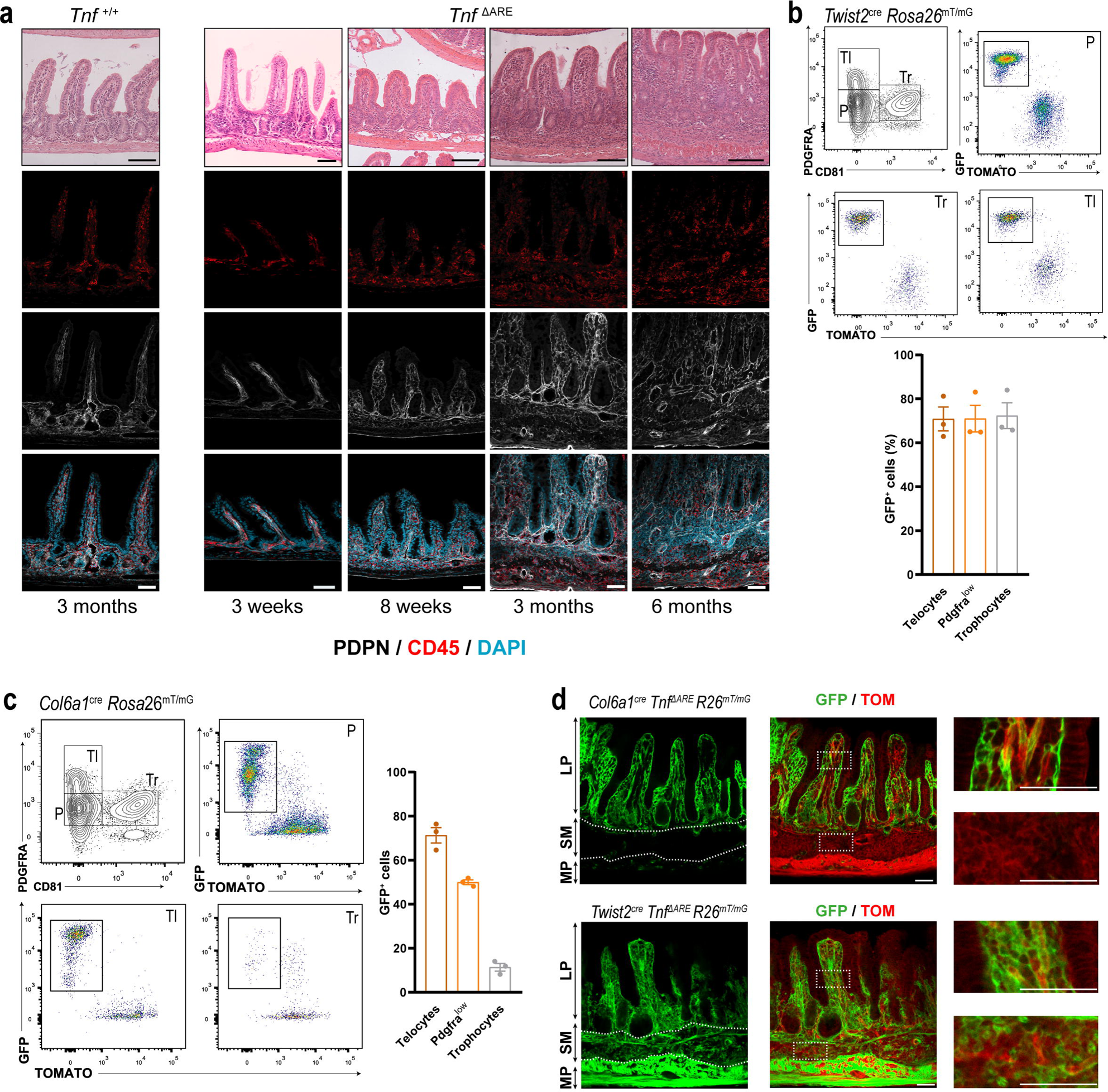
*Col6a1*^cre^ and *Twist2*^cre^ fibroblast-specific targeting in the ileum. a/Representative H&E images of 4μm-thick ileal paraffin sections and immunofluorescent confocal images from 12μm-thick ileal cryosections, stained with DAPI and antibodies against Podoplanin (PDPN) and CD45. b,c/Flow cytometry analysis of Live/Lin^−^/Pdpn^+^ ileal fibroblasts from the ileum of 3 month-old Twist2^cre^ R26^mT/mG^ (b) (*n*=3) and Col6a1^cre^ R26^mT/mG^ (c) (*n*=3) reporter mice. Barplots indicate data ± SEM. Quantification of GFP+Tom-populations is represented as a percentage in the different fibroblast subsets. Tl:Telocytes, P: Pdgfra^low^, Tr: Trophocytes d/Representative confocal images from the ileum of 3-month old *Twist2*^cre^ *R26*^mT/mG^ *Tnf*^ΔΑRE^ and *Col6a1*^cre^ *R26*^mT/mG^ *Tnf*^ΔΑRE^ mice. White dashed boxes indicate areas of higher magnification depicted on the right. LP: lamina propria, SM: submucosa, MP: muscularis propria. Scale bars: 50μm.

We next used two different fibroblast-specific cre lines to target intestinal fibroblasts; the *Twist2*^cre^ and the *Col6a1*^cre^ mice.^26, 27^ Crossing these mice with the *R26*^mTmG^ reporter strain^28^ revealed that *Twist2*^cre^ efficiently targeted a significant proportion of fibroblasts (approximately 65%), while *Col6a1*^cre^ seemed to have a more restricted targeting efficiency (approximately 50%) (SFig. 9a). Upon conducting a more detailed analysis, we found that the *Twist2*^cre^ line exhibited broad specificity across fibroblast subpopulations, including telocytes, trophocytes, and Pdgfra^low^ fibroblasts (Fig. 5b). In contrast, the *Col6a1*^cre^ mice, similarly to published results,^29^ displayed high specificity for telocytes (approximately 70%) and Pdgfra^low^ fibroblasts (approximately 50%), but their targeting efficiency for trophocytes was notably low (approximately 10%)(Fig. 5c).

Given this discrepancy in targeting homeostatic fibroblast subpopulations, we were intrigued to investigate whether these distinct patterns of targeting persisted in the *Tnf*^ΔΑRE^ ileum. Since, telocytes are loosing the high expression of Pdgfra in the *Tnf*^ΔΑRE^ mice (both in mRNA and protein level) (SFig. 9b, SFig. 9c), we decided to investigate the specificity of fibroblast populations through immunofluorescence in the inflamed ileum. During an advanced disease stage (3 months), we found that *Twist2*^cre^-expressing cells were widely distributed throughout the inflammatory lesions, including areas where telocytes are typically found (subepithelial regions), intra-villus space, the submucosal layer, and even within the muscle layer. (Fig. 5d) Interestingly, myocytes also expressed GFP, both in the circular and longitudinal muscle layers (Fig. 5d). On the other hand, fibroblasts targeted by the *Col6a1*^cre^ exhibited a different distribution pattern, being exclusively present in the lamina propria, including subepithelial regions, and inside the villus space Notably, the *Col6a1*^cre^ line did not efficiently target fibroblasts in the submucosal layer (Fig. 5d).

Taken together, these results indicate that the *Twist2*^cre^ strain shows broad specificity towards fibroblasts in both healthy and *Tnf*^ΔΑRE^ ilea. However, the *Col6a1*^cre^ line, which specifically targets telocytes and Pdgfra^low^ subsets in healthy mice, exhibits restricted targeting efficiency in the inflamed lamina propria of *Tnf*^ΔΑRE^ mice.

### 6/Different fibroblast subsets are responsible for triggering and advancing ileal inflammation via TNFR1 signaling

Given the high activation of TNF signalling in *Tnf*^ΔΑRE^ fibroblasts as revealed by our scRNA-seq analysis, we sought to identify the role of *Tnfrsf1a* (gene encoding the TNFR1, the major receptor for TNF) expressed in fibroblasts during ileitis progression.

To delve into this, we generated fibroblast-specific *Tnfrsf1a* deficient *Tnf*^ΔΑRE^ mice using the *Col6a1*^cre^ line. Depletion of *Tnfrsf1a* in telocytes and Pdgfra^low^ cells completely prevented the development of inflammation in the ileum at 3 months of age (Fig. 6a). Even at the chronic disease stage (6 months), the mice only exhibited minimal immune cell infiltration, primarily confined to the submucosal layer (Fig. 6b, 6c). Intriguingly, this was accompanied by an expansion of collagen-producing fibroblasts and myeloid cells in this area (SFig. 10a,b). These findings underscore the critical role of Tnfrsf1a-expressing fibroblasts, especially the telocytes and Pdgfra^low^ subsets, in triggering inflammation in the lamina propria.

**Figure 6.**
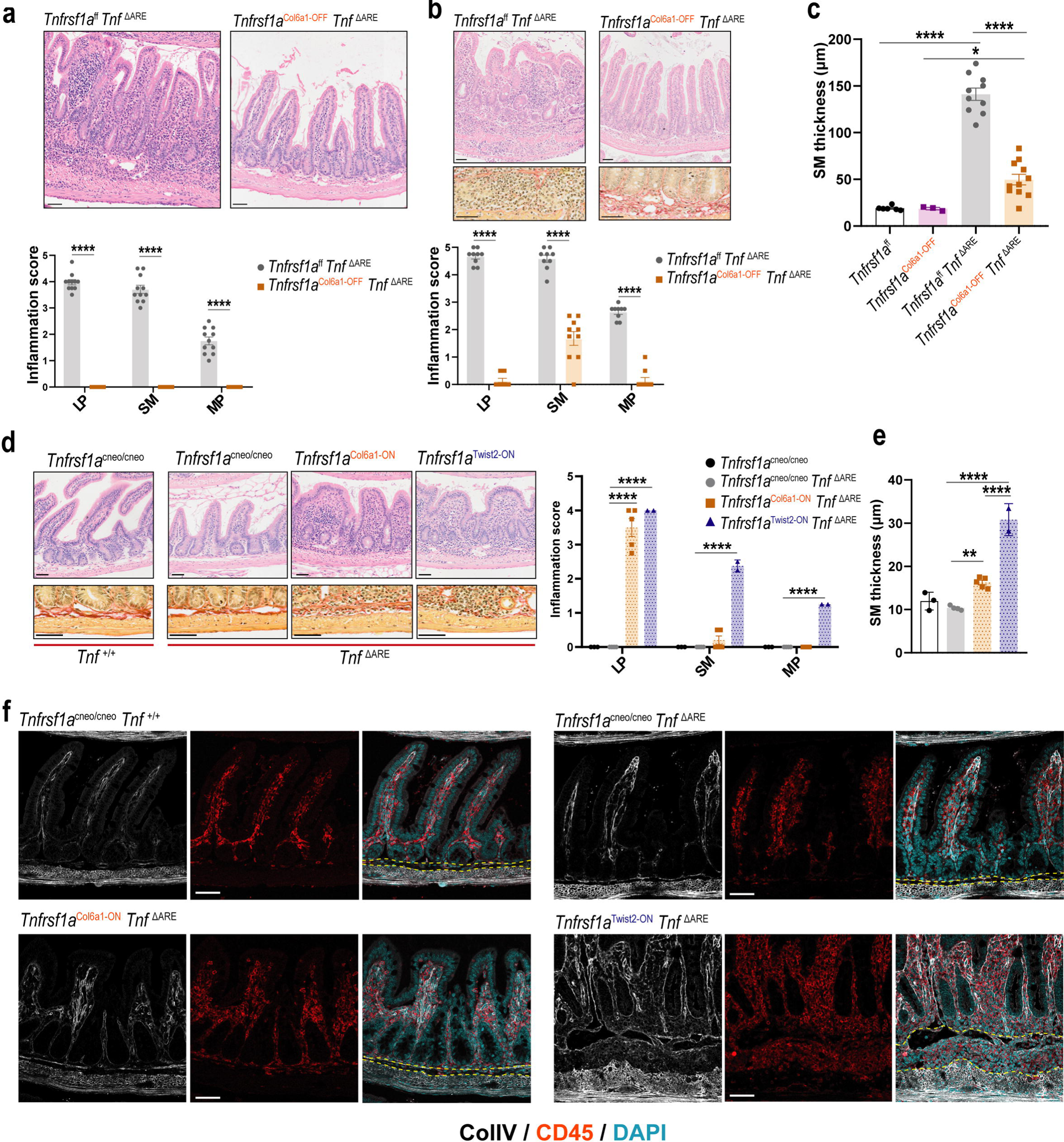
Distinct fibroblast populations orchestrate ileitis development and progression in the different intestinal layers. **a,b/** Representative H&E stained histological images and inflammation scoring of the different intestinal layers in the ileum of 3-month (a) and 6-month (b) old mice with the indicated genotypes. Additionally, representative images of the ileal submucosal layer stained with Sirius Red in 6-month-old mice. (b). c/Quantification of the submucosal thickness (μm). d,e/ (d) Inflammation scoring in the different ileal layers and representative H&E and Sirius Red images of 4-month old mice. (e) Submucosal thickness quantification. (f) Immunofluorescence in paraffin sections from the ileum of 4-month-old mice stained with ColIV, CD45, and DAPI. Yellow dashed lines define the limits of submucosal layer. LP: Lamina propria, SM: Submucosal layer, MP: Muscularis propria. Scale bars: 50μm. one (c,e) or two (a,b) way ANOVA. Each dot represents sample from 1 mouse, *n*=3-11 per genotype, data ± SEM, *p.value < 0.05, **p.value < 0.01, ***p.value < 0.001, ****p.value < 0.001,

To better understand the requirement of *Tnfrsf1a* expression in distinct fibroblast subpopulations, we employed the *Tnfrsf1a*^cneo^ mutant mice, which harbor a conditional gain-of-function allele for TNFR1 receptor. In these mice, *Tnfrsf1a* expression is blocked by a floxed neomycin cassette, but it can be reactivated through Cre-mediated neo excision. *Col6a1*^cre^ driven expression of *Tnfrsf1a* in telocytes and Pdgfra^low^ cells of *Tnf^ΔΑRE^* mice induced inflammation restricted in the lamina propria that did not extend into the deeper intestinal layers (submucosa and muscularis mucosa) (Fig. 6d). In contrast when *Tnfrsf1a* was expressed by a broader category of fibroblasts subsets, including trophocytes, inflammation extended beyond the lamina propria and reached the submucosa and muscle layer (Fig. 6d). Also, it became evident that submucosal thickness increased significantly only when *Tnfrsf1a* was expressed in all fibroblast subsets, not solely in telocytes and Pdgfra^low^ (Fig. 6e). The enlargement of submucosa in *Tnfrsf1a*^Twist2-ON^ *Tnf*^ΔΑRE^ mice was accompanied by immune cell infiltration and fibroblast expansion, indicative of the full development of ileitis in these mice(Fig. 6f).

Overall, these findings suggest that telocytes and/or Pdgfra^low^ cells are required for the initiation of infllammation in the lamina propria. In addition, expression of *Tnfrsf1a* from the same cells suffices for the development of inflammation in the lamina propria. However, only when *Tnfrsf1a* was expressed in all fibroblast subsets (including the trophocytes), there was full expansion and progression of ileitis.

## Discussion

Over recent years, various studies have illuminated the previously underestimated cellular and functional diversity within the intestinal microenvironment during inflammation and under diverse treatment regimens in IBD patients.^11,12,13,25^ However, animal IBD models, have proven to be invaluable tools for unraveling essential disease-associated mechanisms. Particularly, the *Tnf*^ΔΑRE^ mice, a widely utilized model of CD-like ileitis, played an important role in unambiguously establishing TNF’s causal involvement in the inflammatory response of CD.^5,6^ This discovery followed the initial approval of a monoclonal antibody targeting TNF for CD patients.^30,31^ In this context, comprehending the pivotal cellular actors and molecular pathogenic drivers contributing to the development of *Tnf*^ΔΑRE^-associated pathology holds potential for the stratification and treatment of CD patients exhibiting TNF-dependent inflammatory engagement of the ileum.

Our study offers valuable insights into the intricate cellular complexity and variability observed in the *Tnf*^ΔΑRE^ ileum. By conducting a comprehensive single-cell resolution analysis of immune and stromal cells residing in the inflamed ileum, we were able to identify specific changes in population distribution as well as significant transcriptional and functional alterations during the development of the disease with relelevance to human CD-ileitis.

We initially focused on the immune compartment and identified significant transcriptional changes and shifts in cell population abundances within both myeloid and lymphoid cells. Among the various subsets of T-cells, noteworthy modifications in gene expression were observed specifically in memory T-cells and Th17 cells. These changes led to enhanced effector functions, proliferation, and T-cell activation programs in the *Tnf*^ΔΑRE^ populations. While the existence of effector and memory Th17 cells has been recognized in individuals with IBD, our understanding of the gene program changes that occur within these cells during the disease remains limited.^16,32^ Particularly, the role of memory T-cells in IBD has been a subject of debate.^33^ Our findings indicate that memory T-cells adopt similar gene expression profiles as effector cells, suggesting their potential contribution to the pathogenesis of chronic inflammation.

In the myeloid compartment, we observed a substantial expansion of granulocytes. It’s worth noting that many human IBD single-cell studies omit granulocytes^11,12^ due to challenges arising from the low RNA content and rapid RNA degradation during scRNA-seq process. Our mouse data underscore the significance of granulocytes as a primary source of pro-inflammatory chemokines, including Tnf, IL1a, IL1b, and IL23a. Increased infiltration of neutrophils and activation of fibroblasts have previously been correlated with non-response to several treatments through IL-1R signaling in fibroblasts.^23^ Our results indicate that even in a TNF-dependent model, IL1R signaling and neutrophil chemoattractants are significantly enriched in diseased fibroblasts, highlighting the important role of TNF in orchestrating fibroblast activation and neutrophil infiltration.

Focusing on the macrophage populations, we identified two distinct monocyte-derived lineages that are enriched in either *Tnf*^ΔΑRE^ or healthy mice. During steady-state conditions, resident macrophages are more abundant and are engaged in functions related to endosomal processing and antigen presentation. Conversely, in *Tnf*^ΔΑRE^ mice, monocytes predominantly differentiate into activated macrophages with proinflammatory functions, also exhibiting a significant enrichment in ECM-related proteins. Notably, activated macrophages emerge as the primary source of IL-6 family cytokines. The targeting of signaling pathways associated with various IL-6 family ligands holds promise as a therapeutic strategy for IBD, and multiple ongoing clinical trials are exploring this avenue.^34,35^ Additionally, there is evidence of ECM-remodeling gene expression in granuloma-associated macrophages across diverse tissues, suggesting a potential localization of these macrophages within the ileal granulomas of *Tnf*^ΔΑRE^ mice.^18^ Consequently, IL-6 ligand production may play a role in the formation and function of these macrophage structures during ileal inflammation.

One additional notable finding was the evident expansion of B cells compared to other lymphocytes. This expansion occurs within TLOs, major sites where B cells are segregated during inflammation, forming active germinal centers (GCs) supported by FRC networks. Previous studies have also noted increased TLOs in both IBD patients^36^ and *Tnf*^ΔARE^ mice.^37^ However, the precise role of TLOs in the disease remains a subject of debate.^38^ Interestingly, a recent study indicated that depletion of intestinal TLOs led to increased susceptibility to C. rodentium infection.^39^ Nevertheless, in our *Tnf*^ΔARE^ mice, the absence of TLOs and PPs did not impact the progression of ileitis, consistent with genetic depletion of B cells that did not influence Tnf^ΔARE^ pathology.^40^ These findings suggest that the expansion of TLOs and B cells is more likely a consequence of inflammation rather than a contributing factor to the onset and progression of the disease. In addition, the abberant number of autocrine and paracrine interactions developed by the FRC/FDC/MRC subsets reported in our analysis propose pathways propably required for the proper organization and function of these ectopically developed lymphoid structures.

Our single-cell analysis prominently underscores the significant remodeling of fibroblast populations during ileitis. Despite their diminished numbers, likely attributed to the shortened and blunted intestinal villi, these fibroblasts appear to adopt a robust pro-inflammatory phenotype, irrespective of their specific subset distinctions. The expansion of inflammatory fibroblasts has been previously recognized by numerous studies involving IBD patients.^11,12,13,14^ However, these fibroblasts were categorized as a distinct population, not correlating with the homeostatic fibroblast populations. Our analysis demonstrates that the pro-inflammatory fibroblasts associated with IBD represent activated states of the pre-existing fibroblast populations (telocytes, Pdgfra^low^ cells, and trophocytes) in the *Tnf*^ΔΑRE^ ileum. Cell communication analysis revealed a pronounced propensity for fibroblasts to engage in communication within the intestinal microenvironment, acting as both active senders and receivers. They mainly serve as sources of chemoattractants for various immune cell types and exhibit heightened responsiveness to diverse cytokine signals. Notably, among these signals, TNF signaling stands out as significantly activated in inflamed fibroblast populations, mirroring findings from studies involving human CD patients. The major sources of TNF were identified as granulocytes and activated macrophages. Indeed, previous evidence showcasing the exclusive production of TNF by myeloid cells has demonstrated its sufficiency in inducing ileal inflammation.^40^ In a recent study by Thomas et al., myeloid cells were identified as prominent contributors to TNF production during CD.^25^ While this study also highlighted T cells as primary TNF sources, our model did not validate this finding, likely due to the inherent heterogeneity seen in CD patients.

It is widely acknowledged that initial inflammation in patients with CD occurs in the lamina propria and subsequently advances to the submucosa.^41^ This process involves the infiltration of immune cells into the submucosal region along with a localized expansion of fibroblasts in that area.^42^ The *Tnf*^ΔARE^ phenotype serves as a reflection of these specific histological characteristics. We have previously reported the sufficiency of fibroblast-specific TNFR1 to drive intestinal inflammation.^27^ Our current study presents compelling evidence indicating that distinct subsets of fibroblasts, modulated by TNFR1 signaling, play a crucial role in triggering inflammation within the lamina propria and driving its progression into the deeper layers of the mucosa. Telocytes are essential for the infiltration of immune cells within the lamina propria. However, to facilitate the advancement of inflammation into the submucosal region, active TNFR1 signaling in trophocytes and/or a subset of Pdgfra^low^ cells is necessary. We cannot disregard the possibility that *Tnfrsf1a* expressed by muscle cells (targeted by Twist2cre) may also be essential for the development of transmural inflammation during later stages of chronic disease.

Overall, our study provides valuable insights into the complex cellular dynamics driving TNF-dependent ileitis. We highlight the involvement of various cell subsets, cytokine signaling pathways, and fibroblast-mediated interactions that collectively contribute to disease initiation and progression. Different subsets of fibroblasts have been demonstrated to play a pivotal role in the development of localized inflammation through TNFR1 signaling. Consequently, targeting these distinct fibroblast subsets could serve as a strategy to inhibit the onset of inflammation during its early stages and prevent its progression into the chronic phase.

## Material and methods

### Mice

*Tnf*^ΔΑRE^,^5^ *Col6a1*^cre^,^27^ *CMV*^cre^,^43^ *Ltbr*^f/f^,^44^ *Tnfrsf1a*^f/f^,^45^ and *Tnfrsf1a*^cneo^ ^46^ strains were described previously. *Twist2*^cre^,^26^ and *R26*^mT/mG^ ^28^ reporter mice were obtained from Jackson Laboratory. All mice were bred and maintained on a C57BL/6 genetic background in the animal facilities of the Biomedical Sciences Research Center “Alexander Fleming” under specific pathogen-free conditions. Experiments were performed in accordance with all current European and national legislation and were approved by the Institutional Committee of Protocol Evaluation in conjunction with the Veterinary Service Management of the Hellenic Republic Prefecture of Attika. All mice were observed for morbidity and euthanized when needed according to animal welfare.

### Histology-Histological evaluation of inflammation

The preparation of intestinal swiss rolls has been previously described.^47^ Paraffin-embedded mouse ileal swiss rolls (6cm length) were sectioned and stained with haematoxylin and eosin (H&E) and Sirius Red. H&E stained ileal sections were evaluated in a blinded, semiquantitative manner to assess inflammation development across different layers of the intestine. The evaluation was based on the following scale (total: 0-13). Lamina propria: Inflammation scale of 0-4 and expansion scale of 0-1 across the 6cm of ileum. Submucosa: Inflammation scale of 0-4 and expansion scale of 0-1 across the 6cm of ileum. Muscularis propria: Inflammation scale of 0-2 and expansion scale of 0-1 across the 6cm of ileum.

### Measurement of submucosal thickness

Submucosal thickness was measured using OlyVIA (Ver.2.9.1) software in whole swiss roll images acquired with an Olympus Slide Scanner VS200 (20X lens). Measurements were made between the muscularis mucosa and the highest myocytes within the circumferential smooth muscle. 30-60 measurements from different ileal areas were taken and averaged for each mouse to provide a quantitative comparison of submucosal thickness.

### Immunofluoresence/Immunohisctochemistry

Ileum (6cm) was fixed with 4% PFA (O/N). Ileal cryosections of 12 μm thickness were rehydrated in wash buffer (0.1% saponin in PBS) for 15 minutes and blocked in PBS containing 1% albumin for 1h. Sections were incubated with the following primary antibodies: anti-CD45 (AF114, R&D Systems), Biotin-conjugated anti-podoplanin (127403, Biolegend), anti-CD68 (PAS-78996, ThermoFisher) and anti-Pdgfra (AF1062, R&D Systems). Unconjugated antibodies were detected with the following secondary antibodies: Alexa Fluor® 647 goat anti-rabbit (A21244, ThermoFisher) and Alexa Fluor® 647 donkey anti-goat (A21247, ThermoFisher). Biotinylated antibodies were detected using Alexa Fluor® 647 -Streptavidin (S21374, ThermoFisher) and Alexa Fluor® 488-Streptavidin (S32354, ThermoFisher).

For paraffin sections (4 μm thickness), antigen retrieval was performed using Sodium Citrate Buffer (10mM Sodium Citrate, 0.05% Tween 20, pH 6.0) with microwave heating. Tissue permeabilization was achieved with 0.03% Triton in PBS for 10 minutes, followed by blocking in PBS containing 1% albumin for 1 hour. Immunofluorescence staining on these sections involved primary antibodies against anti-CD45 (AF114, R&D Systems), Biotin-conjugated anti-podoplanin (127403, Biolegend), anti-CD68 (PAS-78996, ThermoFisher), and anti-collagen IV (ab6586, Abcam). Secondary antibodies, including Alexa Fluor® 647 goat anti-rabbit (A21244, ThermoFisher), Alexa Fluor® 488 Donkey anti-Rabbit (A21206, ThermoFisher), and Alexa Fluor® 647 donkey anti-goat (A21247, ThermoFisher), were used for signal detection.

For B220 staining, Biotin-conjugated anti-B220 (103204, Biolegend) was applied, and signal detection and amplification were performed using the ABC kit (Vector Laboratories) with Vectastain DAB (3,3-diaminobenzidine) kit (Vector Laboratories) for signal development, followed by hematoxylin counterstaining.

Single cell suspensions from the ileum were concentrated onto slides using a Cytospin™ Centrifuge. The slides were fixed with methanol, permeabilized with 0.1% saponin in PBS, and blocked with PBS containing 1% albumin. They were then incubated with primary antibodies: eFluor™ 660 anti-CD68 (50-0681-82, eBioscience) and anti-Lysozyme EC 3.2.1.17 (A0099, Dako). The secondary antibody used for the detection of Lysozyme was Alexa Fluor® 488 goat anti-rabbit (A11008, ThermoFisher).

Stained sections were mounted with Fluoroshield with DAPI (F6057, Sigma). Imaging was performed using a TCS SP8X White Light Laser confocal system (Leica), a Zeiss LSM900 confocal microscope, and an Olympus Slide Scanner VS200.

### Flow cytometry

The isolation of cells from small intestine lamina propria was performed as previously described.^29^ In summary, the terminal ileum (6cm length) was prepared by removing Peyer’s patches. The intestine was longitudinally opened and subjected to a 30-minute incubation at 37°C in HBSS (14170-088, Gibco) containing 5 mM EDTA, DTT (D9779, Sigma), and 10 mM Hepes (LM-S2030, Biosera). After thorough agitation and PBS washes to eliminate epithelial cells, the remaining tissue underwent digestion using 300 U/ml Collagenase XI (C7657, Sigma), 0.8 u/ml Dispase II (18538700, Roche), and 100 U/ml Dnase I (DN25, Sigma) for 40–60 minutes at 37°C. The resulting cell suspension was filtered through a 70 μm strainer, followed by centrifugation and resuspension in FACS buffer (PBS with 2% FBS). For stainings, 1–2 million cells/100 μl were incubated with the following antibodies: anti-CD11c PeCy7 (117318, Biolegend), anti-CD11b (101212, Biolegend), anti-CD45 Alexa Fluor 700 (103128, Biolegend), anti-EPCAM APC-Cy7 (118218, Biolegend), anti-Podoplanin PeCy7 (127412, Biolegend), anti-CD31 PerCP/Cyanine5.5 (102420 Biolegend), anti-B220 FITC (103206, Biolegend), anti-IgA PE (12420483, Invitrogen), anti-TCRβ FITC (11-5961-85, eBioscience), anti-CD81 Biotin, (13-0811-81, eBioscience), anti-Pdgfra BV605 (135916, Biolegend). As a secondary staining for anti-CD81, Alexa Fluor® 647 -Streptavidin (S21374, ThermoFisher) was used. For intracellular staining (against IgA) cells were fixed and permeabilized using the Fixation and Permeabilization Buffer Set (88-8824-00, eBioscience), according to manufacturer’s instructions. Propidium Iodide (P1304MP, Sigma), DAPI (D1306, Invitrogen) or the Zombie-NIR Fixable Viability Kit (423105, Biolegend) was used for live-dead cell discrimination. Absolute number quantification was performed using Precision Count Beads (424902, Biolegend). Samples were analyzed using the FACSCanto II flow cytometer (BD) or the FACSAria III cell sorter (BD) and the FACSDiva (BD) or FlowJo software (FlowJo, LLC).

### ScRNA-seq library preparation and sequencing

FACs sorted intestinal cell populations from the terminal ileum of 3-month-old *Tnf*^+/+^ and *Tnf*^ΔΑRE^ mice were mixed to an equal ratio and subjected to 10X Chromium Single Cell 3’ Solution v3.1. The scRNA-seq libraries were prepared following the 10X Genomics user guide Single Cell 3’ v3.1 reagent kits pooled and sequenced with the DNBSEQ-G400 sequencer (PE100) (BGI Genomics). The original image data was converted to sequence data via base calling, leading to the storage of raw reads in FASTQ file format. Reads (1,207,791,679 in *Tnf*^+/+^ sample and 776,706,111 in *Tnf*^ΔΑRE^ sample) were then aligned to the mouse reference genome (mm10). The steps of read alignment and gene count summarization per cell, were performed using the 10X Genomics Cell Ranger pipeline (v 7.0.1), resulting in 2,292 and 2,639 median genes detected per cell in *Tnf*^+/+^ and *Tnf*^ΔΑRE^ sample respectively.

### Sequencing data processing for mouse data

Using the filtered output of cellRanger, we performed a first round of doublet detection running scrublet (v 0.2.3) with default settings in both *Tnf*^ΔΑRE^ (25,771 cells) and *Tnf*^+/+^ (43,119 cells) samples. This resulted in the removal of 3,063 and 7,232 predicted doublets respectively. Next, each sample was processed separately. For both samples low quality cells (genes detected or >7,500 and/or percentage of reads mapped to mitochondrial genome >5%) were discarded. Additionally, log-normalization, highly variable genes detection (MVP), scaling of the normalized data and PCA analysis (*Tnf*^+/+^ nPCs: 20, *Tnf*^ΔΑRE^ nPCs: 22) were executed. Regarding clustering, the Louvain algorithm was employed with a low resolution (res = 0.01) in order to separate the cells in the three major compartments (stroma, lymphoid, myeloid). Following that, three separate objects (one for each compartment) was stored for each sample and subjected to a second round of doublet detection analysis utilizing Doublet Finder package (v 2.0.3) with a doubletRate of 0.046, this resulted in the removal of 1,141 WT and 942 DARE cells predicted as doublets. After, integration (Seurat default settings: VST, CCA) was performed for stroma cells (nPCs=23, res=0.5), myeloid (nPCs=20, res=0.5) and lymphoid (nPCs=15, res=0.5) cells between *Tnf*^+/+^ and *Tnf*^ΔΑRE^ samples. Cluster annotation was based on the marker genes, which were calculated using the FindAllMarkers function (pvalue < 0.01, min.pct = 0.25, logfc.threshold = 0.25, Wilcoxon rank sum test) from Seurat package (v 4.3.0). In the stroma compartment, clusters 10, 14 and 19 were excluded from downstream analysis, due to exhibiting both fibroblasts and immune markers (Ptprc, S100a9, S100s8). Differential expression analysis between diseased and healthy conditions was performed using the FindMarkers functions (same cut offs as in findAllMarkers and min.pct=0.1). All the above steps of the analysis (unless mentioned otherwise) were also conducted through the Seurat package. Finally, to showcase the differential abundance of cell clusters between samples we used contour plots through the package scDataViz (v 1.6.0).

### Pathway analysis

Functional enrichment analysis and pathway activity at cluster level was achieved using metascape (https://metascape.org/) and PROGENy (v 1.18.0) & decoupleR (v 2.2.2) respectively. Metascape analysis: for stroma, myeloid and lymphoid compartments intra-cluster (*Tnf*^ΔΑRE^ vs *Tnf*^+/+^) up/down regulated genes were used as an input. For the lymphoid compartment positive marker genes were also used to find enriched terms in the different clusters. Additionally, a background genelist was used, considering as active genes all the genes expressed in more than 3 cells. As regards the pathway activity analysis, decoupleR proposed methodology for PROGENy was adopted, as described in the vignette (mouse, top 100 genes per pathway), for *Tnf*^+/+^ and *Tnf*^ΔΑRE^ samples. Activity scores per cell were summarized at cluster level (mean activity).

### Trajectory analysis

In the myeloid compartment, trajectory analysis was performed utilizing Slingshot package (v 2.4.0). More particularly, umap coordinates and cluster labels were used as input, while the root cluster was set to Monocytes. The aforementioned analysis was run in both samples, in each case the two identified lineages were reported.

### Correlation with human datasets

Human ileitis raw counts data and metadata tables were downloaded from https://singlecell.broadinstitute.org/single_cell/study/SCP1884/human-cd-atlas-study-between-colon-and-terminal-ileum. Cells from terminal ileum were selected and grouped in stroma and lymphoid Seurat objects. Default Seurat analysis was employed; however, the original cell type annotation was kept. After conversion of the genes from human to mouse, integration between human and mouse datasets was performed for lymphoid and stroma compartments, following the default settings in Seurat. Next, spearman correlation analysis was performed between the different clusters of *Tnf*^ΔΑRE^ and human inflamed samples using the top highly variable genes. For the myeloid compartment, two gene signatures containing 80 genes were retrieved from a previously published dataset (STab. 4).^11^ Signature scoring was performed using UCell (v 2.0.1), cell scores were summarized at cluster level and plotted in violin plots. For the enrichment analysis presented in Fig. 3h, cells were organized into superclusters following the scheme explained in STab. 5). After differential expression analysis was performed in both human (Inflamed vs Healthy) and mouse fibroblasts (*Tnf*^ΔΑRE^ vs *Tnf*^+/+^). Up and down regulated genes were identified utilizing the findMarkers function from Seurat with thresholds: (pvalue < 0.05, min.pct = 0.1, logfc.threshold = 0.25, Wilcoxon rank sum test). Subsequently, up and down regulated lists of genes were given as input to metascape.

### Cell-Chat

Cell-cell communication analysis was performed with CellChat package (v 1.5.0) for both *Tnf*^+/+^ and *Tnf*^ΔΑRE^ samples using default options. Glial cells were removed prior to analysis, due to the low number of cells. Additionally, the produced cellchat objects were subjected to comparative analysis using default settings. In order to compare the activation of signalling pathways between the two conditions, netP tables from cellchat objects were utilized and the aggregated probability scores were calculated in two scenarios. In the first non-FRCs fibroblasts (trophocytes, telocytes, Pdgfralow) were used as senders and in the second one as receivers. Since communication probability in Cellchat signifies the interaction strength, we calculated a log2 ratio of communication probabilities for all the pathways, which were present in both samples and visualized them in a dotplot format. For the signalling pathways that were found uniquely either in *Tnf*^+/+^ or *Tnf*^ΔΑRE^ we reported the aggregated communication propabilty.

### Statistical analysis

Data are presented as mean±SEM. Student’s t-test (parametric, unpaired, two-sided) one-or two-way ANOVA were used for evaluation of statistical significance using GraphPad 6 software. Statistical significance is presented as follows: * p < 0.05, ** p < 0.01, *** p < 0.001, **** p < 0.0001.

---

**<u>Supplementary Figure 1.</u>** a/Representative H&E stained images from the ileum of 3-month old *Tnf*^+/+^ and *Tnf*^ΔΑRE^ mice. Scale bar=100μm. b/Gating strategy for sorting stroma, myeloid and lymphoid cells for scRNA-seq. c/Cellular distribution of stroma, myeloid and lymphoid cells acquired before sorting and recovered after scRNA-seq.

**<u>Supplementary Figure 2.</u>** a/ Dotplot showing the normalized expression of top marker genes for each lymphoid cell cluster. b/ Absolute number of lymphocytes assessed by flow cytometry and normalized to the length of the ileum in 3-month-old mice of the indicated genotypes. TCRαβ+ Τ cells were gated as CD45^+^TCRβ^+^, Β-cells as CD45^+^B220^+^ cells and IgA^+^ plasma cells as CD45^+^B220^−^IgA^+^. Each dot represents a sample from an individual mouse, data ± SEM, *n* = 6–9. ***p < 0.001, ****p < 0.001, (two-tailed Student’s t test).

**<u>Supplementary Figure 3.</u>** a/ Dotplot showing the normalized expression of top marker genes for each myeloid cell cluster. b/ Barplots depicting differences in the distribution of *Tnf*^+/+^ (green) and *Tnf*^ΔΑRE^ (red) cells along pseudotime in the two different lineages identified by Slingshot. In both cases, pseudotime values are divided in 5 bins. c/ Dotplot representing the normalized expression of selected ECM related genes by myeloid clusters. d/ Representative immunofluorescence image of CD68^+^ macrophages (green) in *R26*^mT/mG^ *Tnf*^+/+^ and *R26*^mT/mG^ *Tnf*^ΔΑRE^ ileum. White staining represents the expression of *Tomato.* Scale bar=50μm e/ Heatmap showing scaled expression at a single cell level of the human resident and proinflammatory markers (STab. 4) identified in the ileum of CD patients.

**<u>Supplementary Figure 4.</u>** a/ Dotplot displaying the normalized expression of top marker genes for each stromal cell cluster. b/ Barplot showing the differential abundance of myeloid cell clusters per genotype. c/Gating FACs strategy for the quantification of the absolute number and percentage of blood endothelial cells (BECs), lymphatic endothelial cells (LECs) and fibroblasts (FB) in the total of live CD45^−^EPCAM^−^ population (stroma).

**<u>Supplementary Figure 5.</u>** a/ Total number of identified cellular interactions and interaction strength in *Tnf*^+/+^ and *Tnf*^ΔΑRE^ cells, calculated by CellChat. b/ Violin plots showing the normalized expression levels of *Tnf, Tnfrsf1a,* and *Tnfrsf1b* in *Tnf*^+/+^ and *Tnf*^ΔΑRE^ cell subsets.

**<u>Supplementary Figure 6.</u>** a-c/ Chord diagram depicting detected signalling pathways using CellChat between FDCs/MRCs (a), FRC1 (b), FRC2 (c) and the rest of the cell clusters in healthy (left panel) and disease state (right panel).

**<u>Supplementary Figure 7.</u>** a/ Chord diagram showing detected signalling pathways from FDCs/MRCs FRC1 and FRC2 to FDCs/MRCs FRC1 and FRC2 in healthy (left panel) and disease state (right panel).

**<u>Supplementary Figure 8.</u>** a/Representative images of ileal swiss rolls stained with anti-B220 and quantification of B220^+^ clusters as TLO structures (PPs were excluded). Each dot represents a mouse with the indicated genotype at the age of 3 months. Quantification was performed using two paraffin sections from different levels and normalized to the total ileum length. b/ Representative H&E images and quantification of inflammation score in 3-month old *Ltbr*^f/f^ and *Ltbr*^KO^*Tnf*^ΔΑRE^ mice. mean ± SEM *,n* = 6–7. ***p < 0.001, ns=not significant (two-tailed Student’s t test).

**<u>Supplementary Figure 9.</u>** a/ Gating strategy to evaluate cell targeting of *Col6a1*^cre^ and *Twist2*^cre^ *R26*^mT/mG^ lines in the ileum of 3-month old mice. Targeting specificity was assessed by the percentage of GFP^+^Tom^−^ cells in the total fibroblast (live Lin^−^Pdpn^+^ cells) population. b/Violin plot showing the average normalized expression of *Pdgfra* in ileal stromal cell subsets from *Tnf*^+/+^ and *Tnf*^ΔΑRE^ mice. c/ FACs plots showing the protein expression of PDGFRA and CD81 in ileal fibroblasts (gated as in a) from 3-month old *Tnf*^+/+^ and *Tnf*^ΔΑRE^ mice.

**<u>Supplementary Figure 10.</u>** a,b/ Representative immunofluorescence images in cryosections of *Tnf*^+/+^ and *Tnf*^ΔΑRE^ ilea stained with antibodies against collagen IV (ColIV), CD45, Podoplanin (PDPN), CD68 and DAPI.

## Author’s Contributions

LI, AP, CT and GK designed the study and interpreted the experimented results. LI, CT, and AP performed the experiments and data analysis. FR performed 10X library preparation. VK contributed to data interpretation. LI wrote the manuscript and prepared the figures with contribution from CT. GK edited and finalized the manuscript. All authors were involved in critically revising the final manuscript.

## Acknowledgements

The authors would like to thank Meropi Gennadi, Anna Katevaini and Michalis Meletiou for their excellent technical assistance, and Sofia Grammenoudi for her help in the FACs sorting. We would also like to thank Fleming’s Animal House, Flow Cytometry and Genomics Facilities.

