## Supplementary files for "Cellular complexity and crosstalk in murine TNF-dependent ileitis: Different fibroblast subsets propel spatially defined ileal inflammation through TNFR1 signalling"

**Supplementary Figure 1**

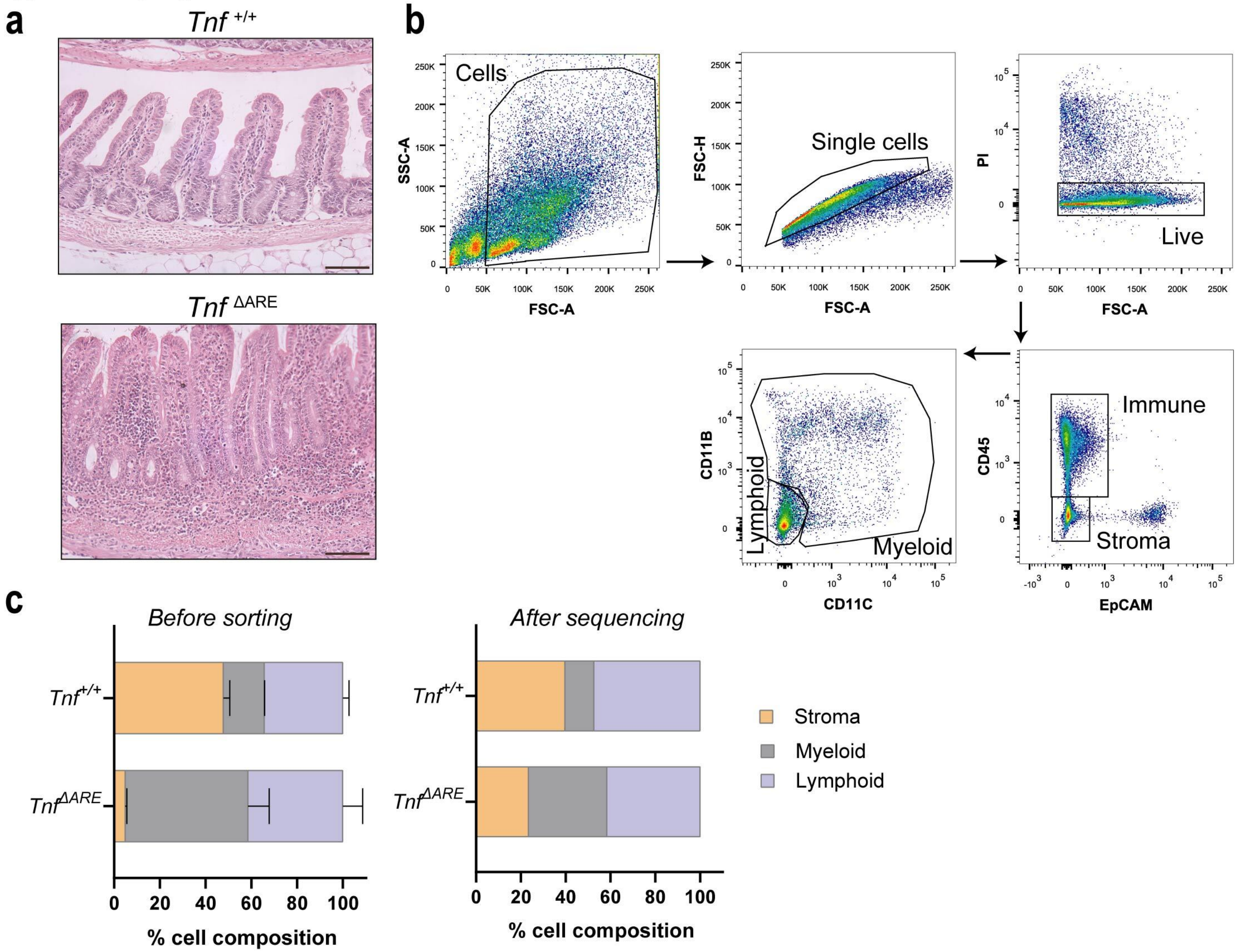

### Supplementary Figure 2

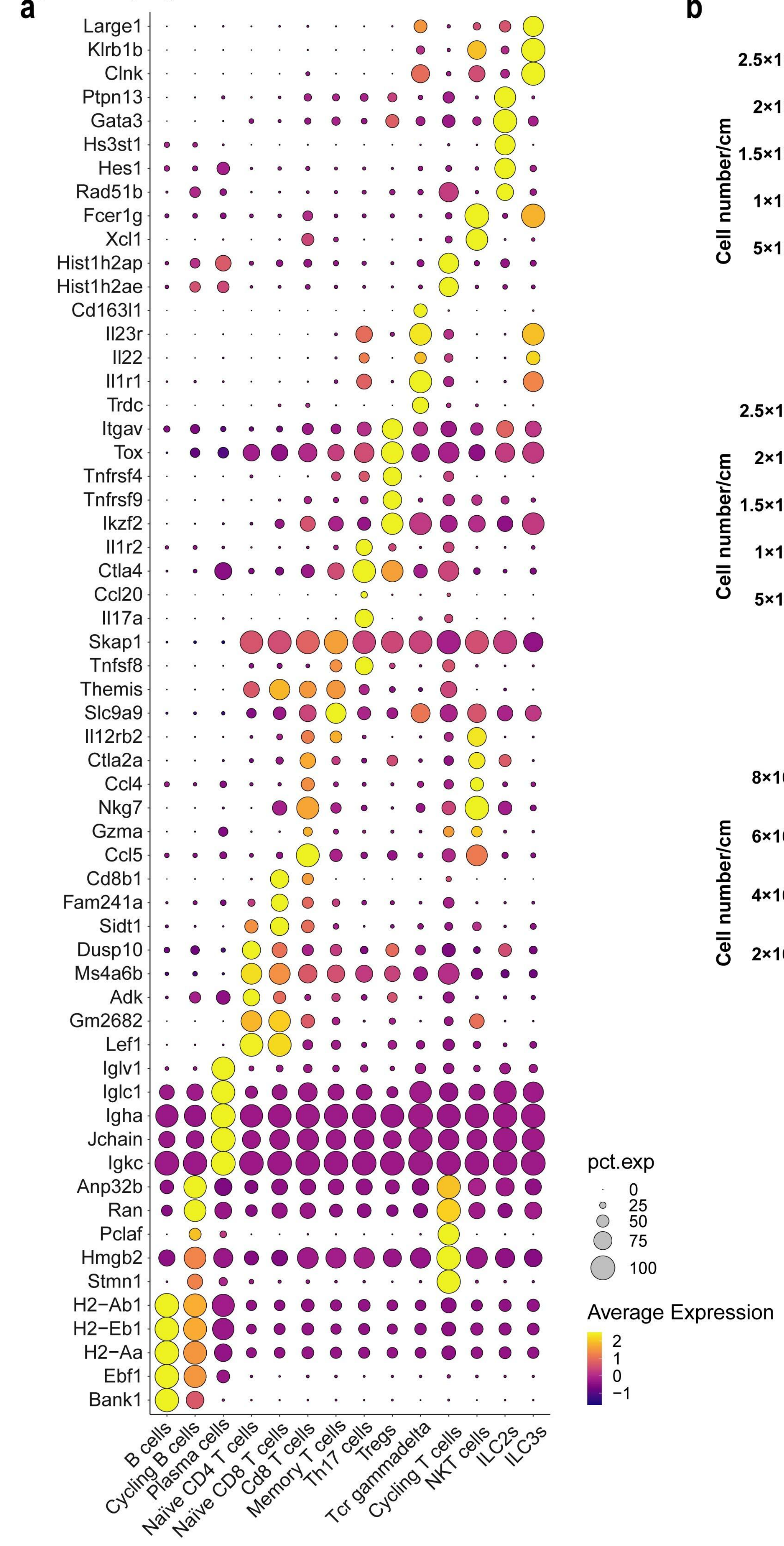

**Supplementary Figure 3**

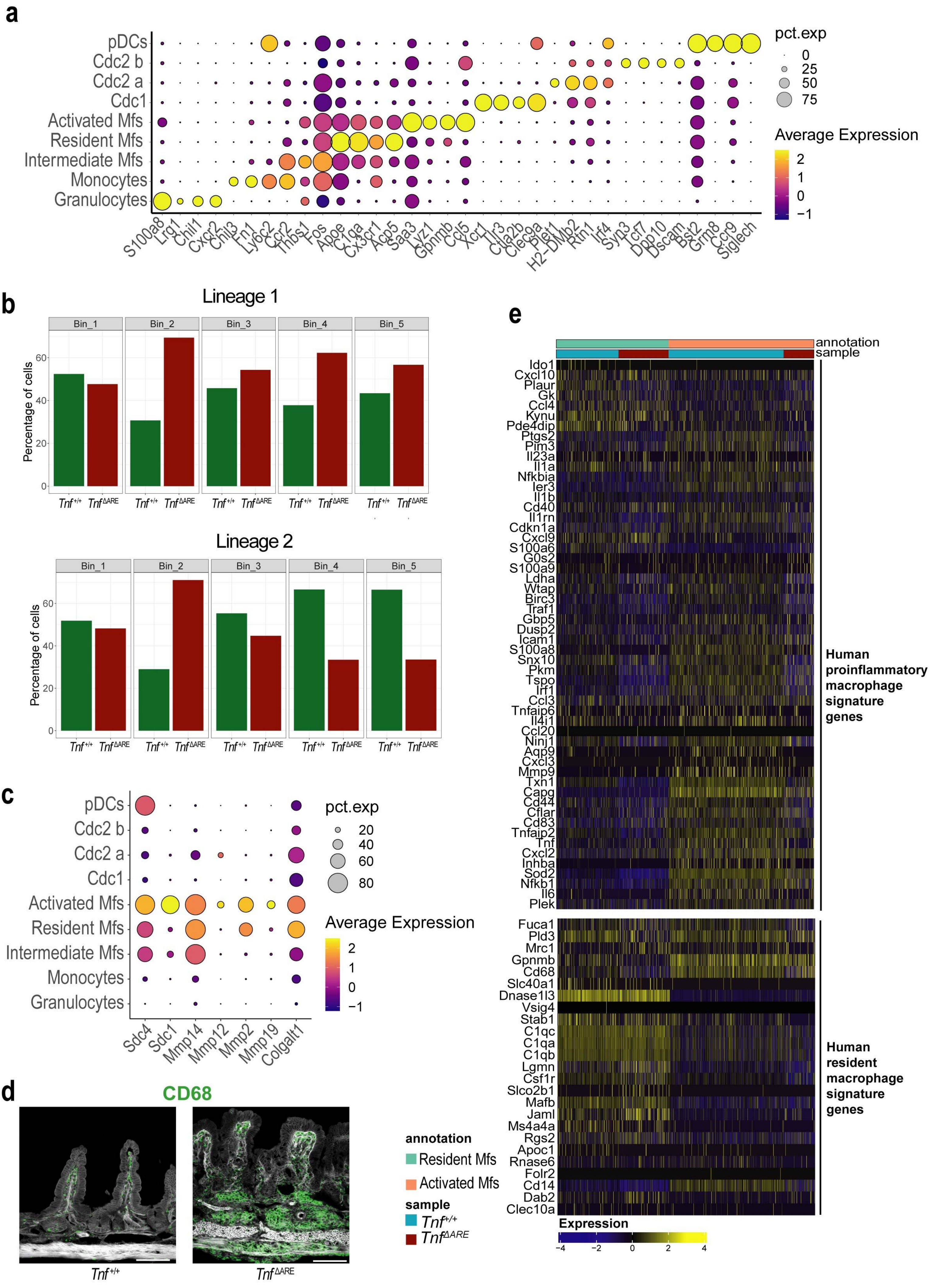

#### Supplementary Figure 4

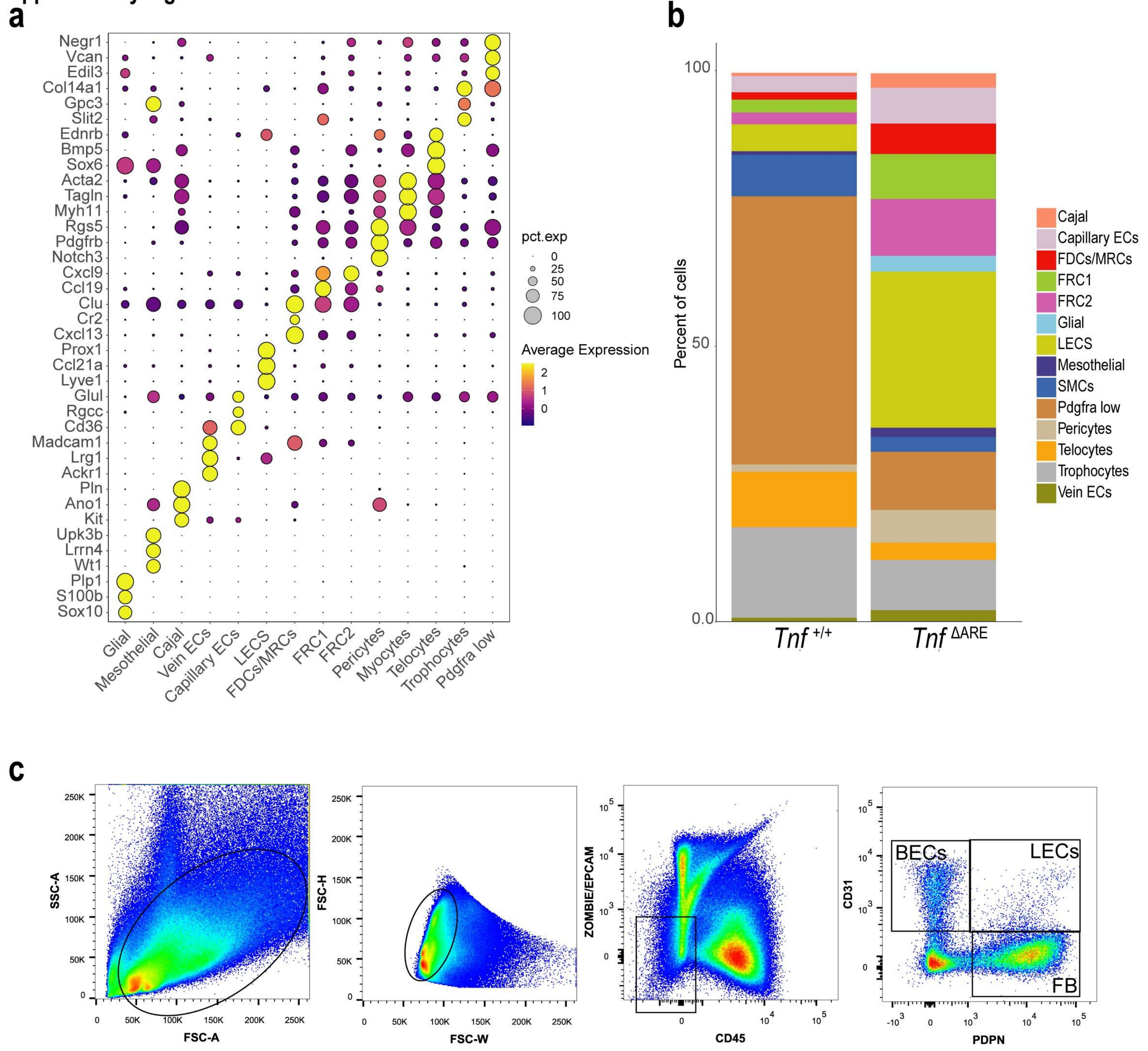

**Supplementary Figure 5**

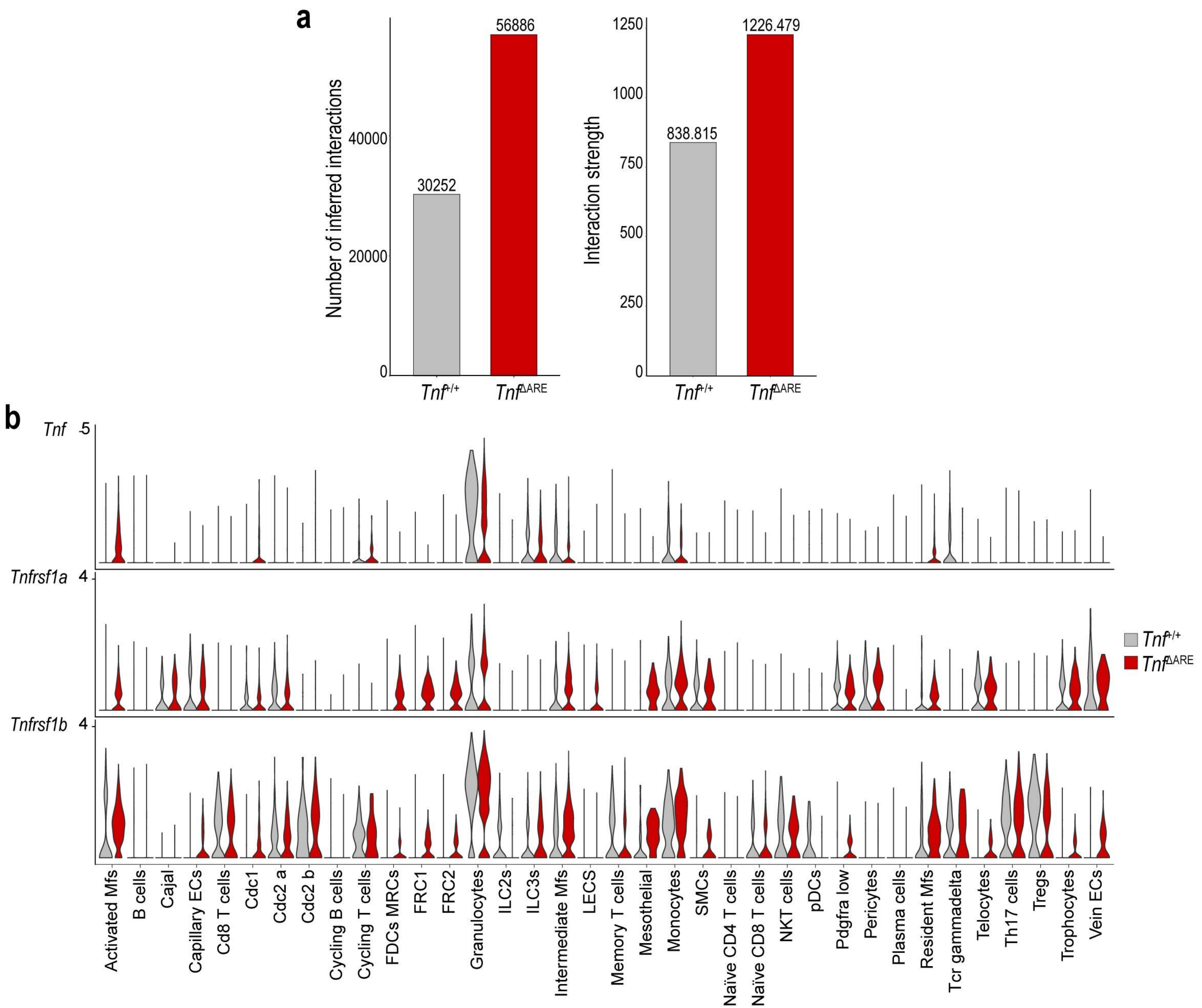

#### Supplementary Figure 6

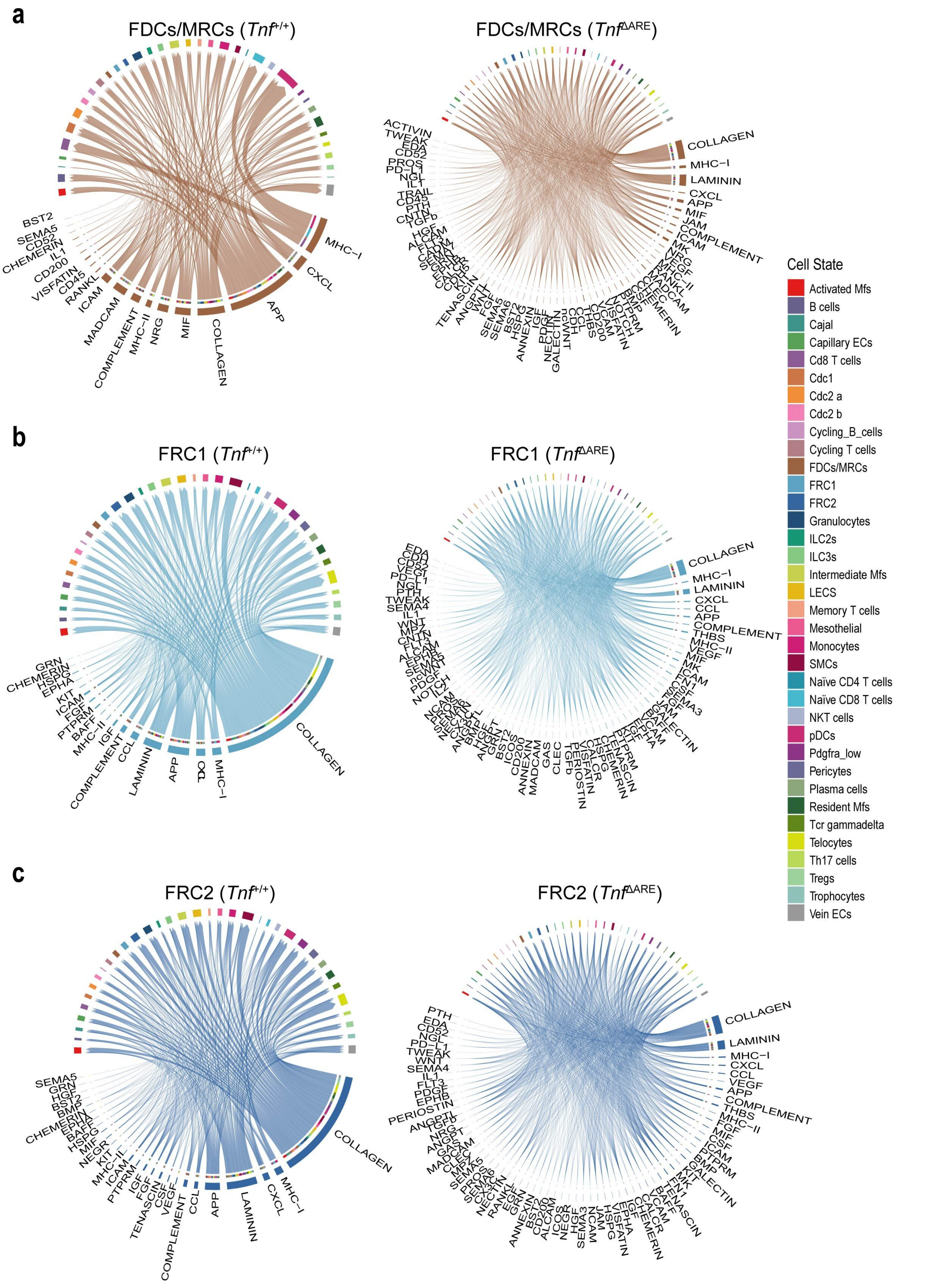

**Supplementary Figure 7**

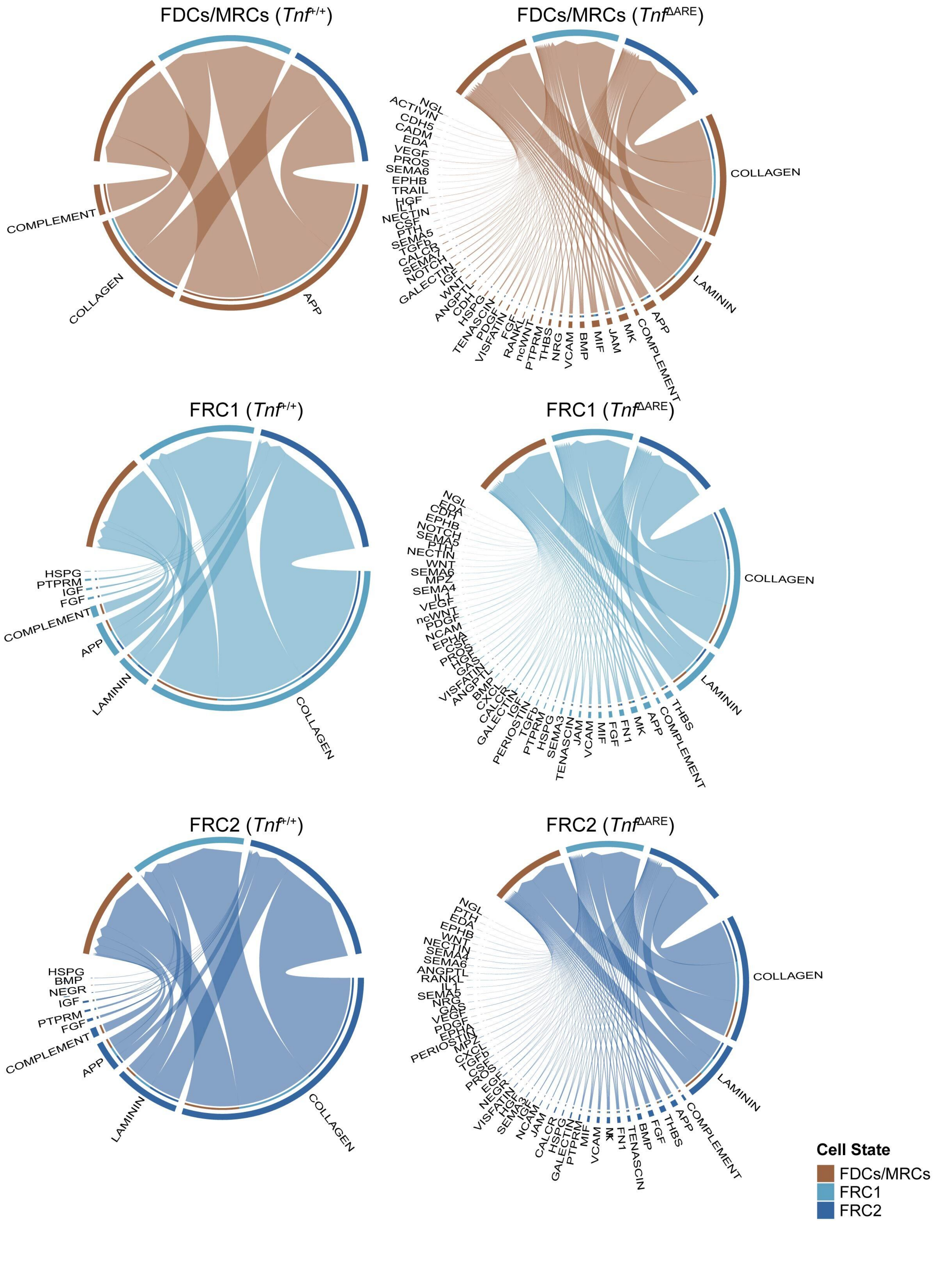

#### Supplementary Figure 8

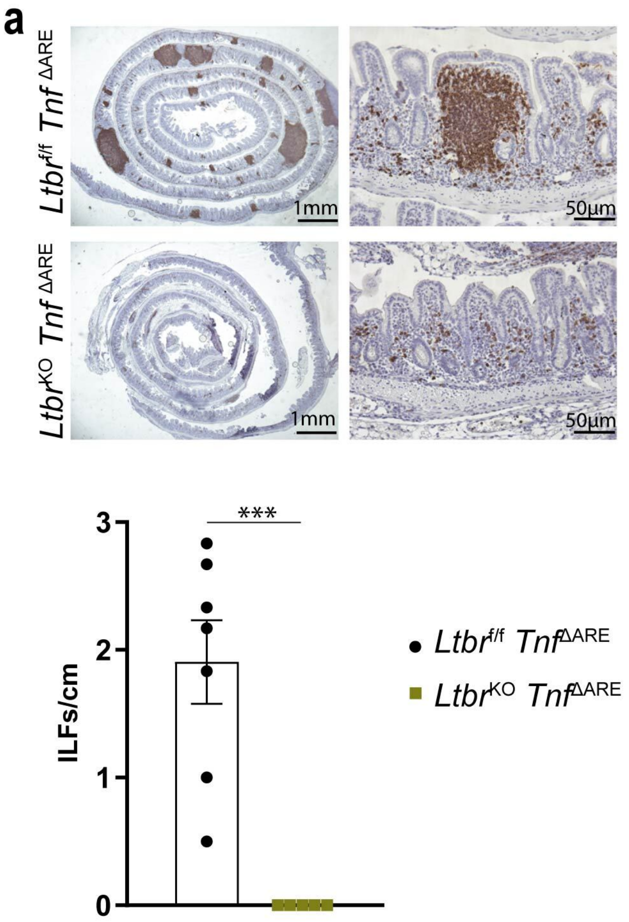


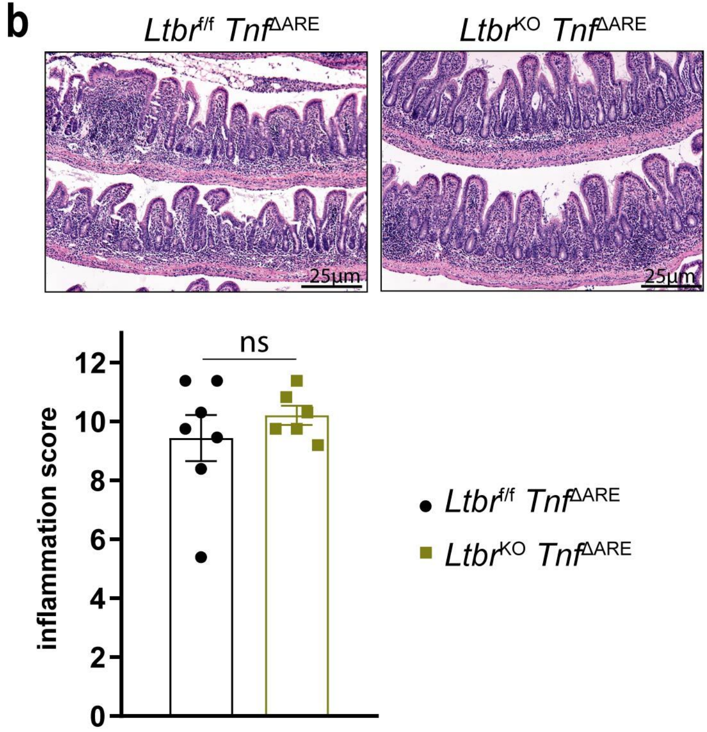

#### Supplementary Figure 9

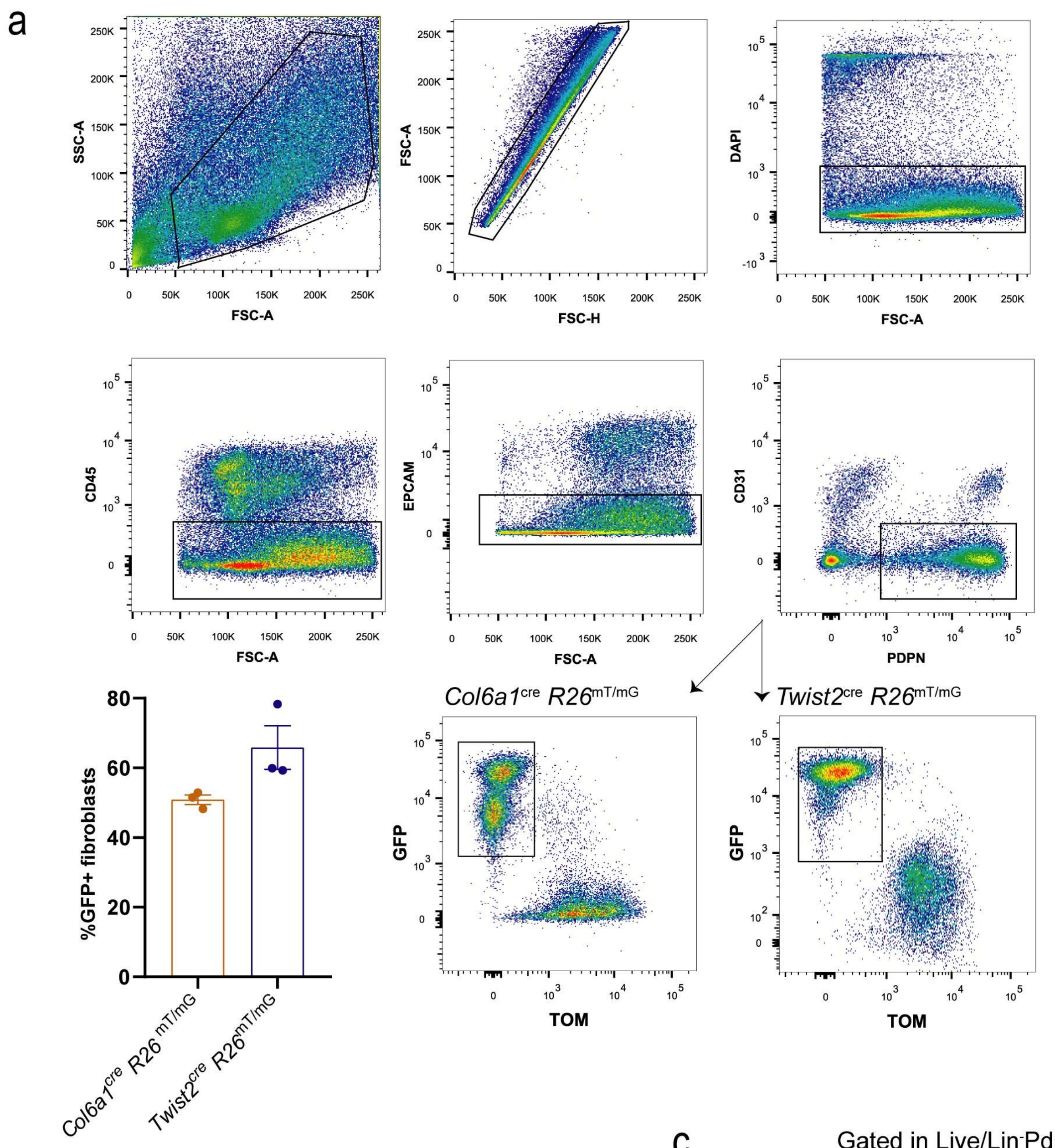


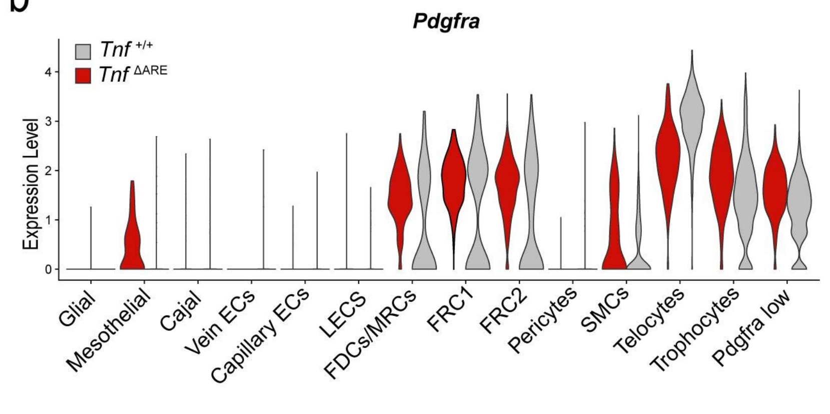


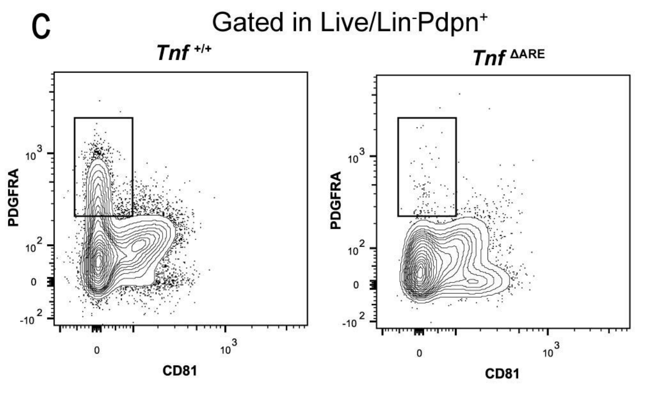

**Supplementary Figure 10**

**a**

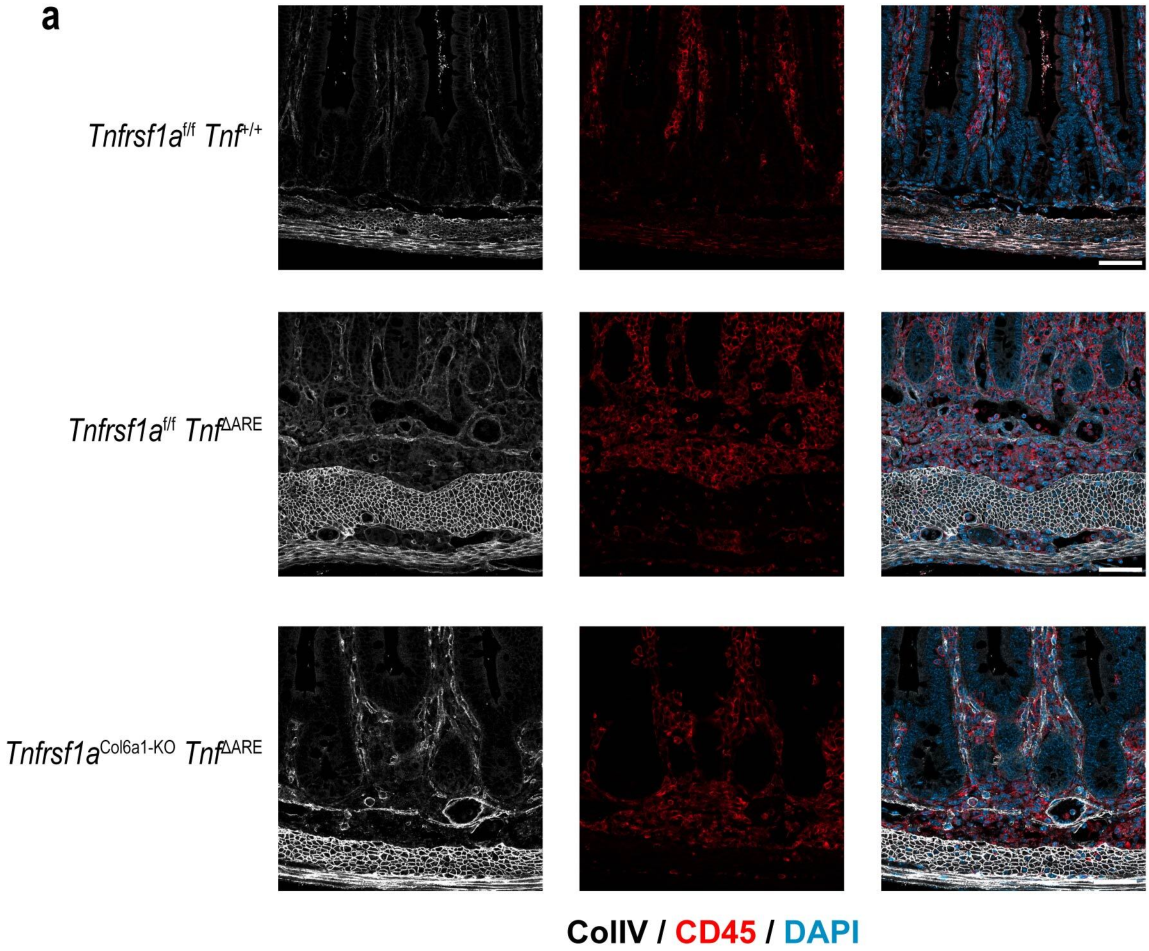


**b**

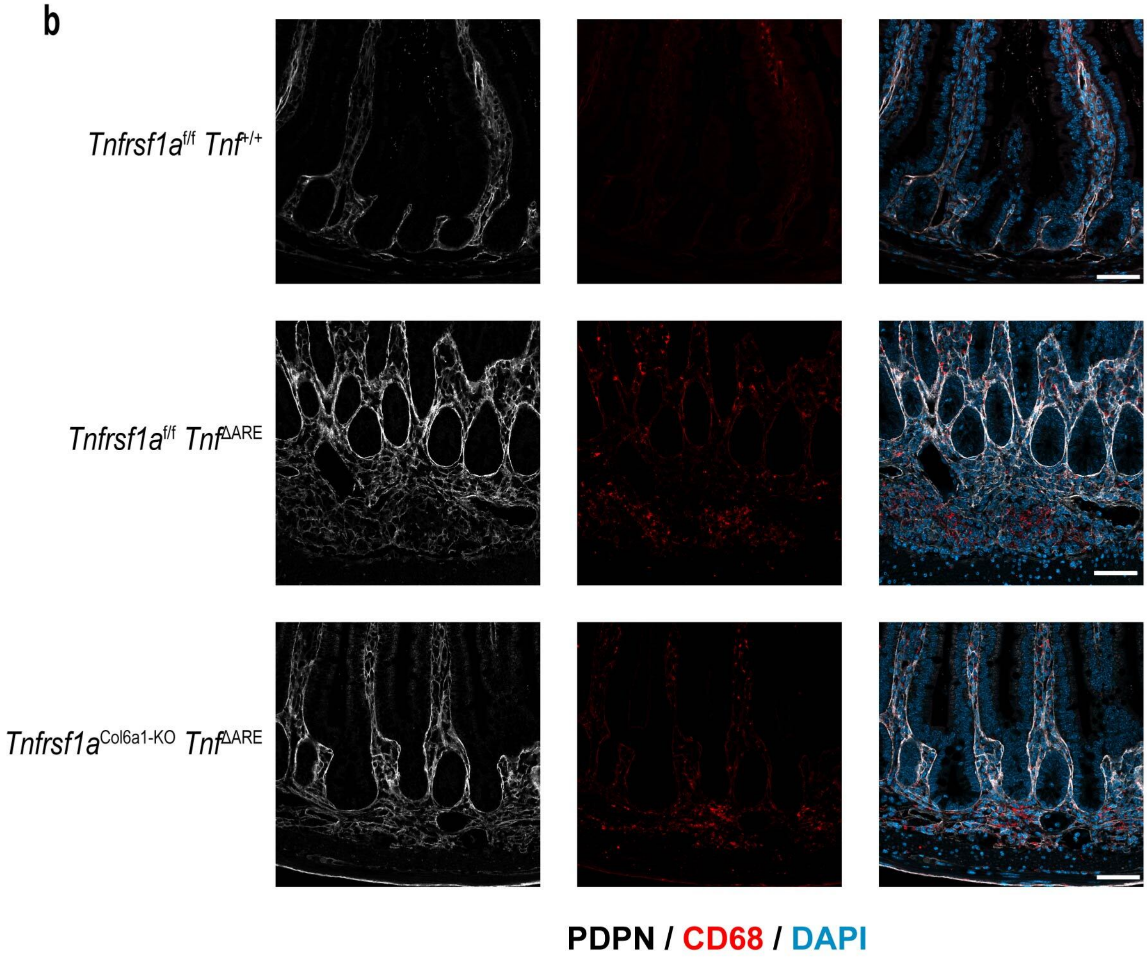
